# Patch repair protects cells from the small pore-forming toxin aerolysin

**DOI:** 10.1101/2022.09.22.509098

**Authors:** Roshan Thapa, Peter A. Keyel

## Abstract

Small pore-forming toxins in the aerolysin family lyse cells by damaging the membrane, but membrane repair responses used to resist them, if any, remain controversial. Four membrane repair mechanisms have been proposed: toxin removal by caveolar endocytosis, clogging by annexins, microvesicle shedding catalyzed by MEK, and patch repair. Which of these repair mechanisms aerolysin triggers is unknown. Furthermore, Ca^2+^ flux triggered by aerolysin is controversial, yet membrane repair responses require Ca^2+^. Here, we determined Ca^2+^ influx and repair mechanisms activated by aerolysin. In contrast to cholesterol-dependent cytolysins (CDCs), removal of extracellular Ca^2+^ protected cells from aerolysin. Aerolysin triggered sustained Ca^2+^ influx. Since aerolysin triggered Ca^2+^ flux, we investigated Ca^2+^-dependent repair pathways. Caveolar endocytosis failed to protect cells from aerolysin or CDCs. MEK-dependent repair did not protect against aerolysin. Aerolysin triggered slower annexin A6 membrane recruitment compared to CDCs. In contrast to CDCs, expression of the patch repair protein dysferlin potently protected cells from aerolysin. We propose that aerolysin triggers a Ca^2+^-dependent death mechanism that obscures repair responses, and the primary repair mechanism used to resist aerolysin is patch repair. We conclude that different classes of bacterial toxins trigger distinct repair mechanisms.

## Introduction

The virulence of many lethal bacteria is mediated partly by the secretion of pore-forming toxins (PFTs). PFTs are the largest family of bacterial toxins, and are classified into subfamilies based on structure. Two subfamilies include the cholesterol-dependent cytolysins (CDCs), secreted primarily by Gram positive bacteria, and the aerolysin family, secreted by a wide range of bacteria. Archetypal toxins representing CDCs include streptolysin O (SLO) from *Streptococcus pyogenes*, and intermedilysin (ILY) from *S. intermedius*. CDCs form large (20-30 nm) pores that require cholesterol for pore formation (Thapa et al., 2020). SLO also uses cholesterol for binding, while ILY binds to the glycosylphosphatidylinositol (GPI)-anchored protein human CD59 (Giddings et al., 2004). Aerolysin is the archetypal aerolysin family member, secreted by *Aeromonas hydrophila*. Aerolysin forms small (4 nm) pores and binds to GPI-anchors (Abrami et al., 1998b). Aerolysin is produced as the inactive precursor pro-aerolysin, which requires cleavage of a C-terminal peptide for activation (Abrami et al., 1998a; Iacovache et al., 2011). Cleavage can occur by cellular furins present on most mammalian cell membranes or *in vitro* using trypsin (Abrami et al., 1998a; Iacovache et al., 2011). After binding, both aerolysin and CDCs form pre-pores on the host plasma membrane that insert and cause cell death if not resisted by the host cell.

Cellular resistance mechanisms are best described for resisting CDCs. To resist CDCs, four Ca^2+^-dependent membrane repair mechanisms have been proposed: toxin removal by caveolar endocytosis, pore clogging by annexins, pore removal by microvesicle shedding, and membrane resealing by patch repair. While caveolar endocytosis was proposed to remove CDCs (Corrotte et al., 2020), no work to date demonstrated removal of intact pores by this mechanism. Instead, we showed that endocytosis acts after repair is complete to remove toxin oligomers and damaged membranes (Romero et al., 2017). In contrast, annexins form a crystalline lattice to physically blockade the pore (Jimenez and Perez, 2017). CDCs are also sequestered on microvesicles, which are subsequently shed (Keyel et al., 2011). While shedding can occur independently of proteins, both the endosomal sorting complex required for transport (ESCRT) (Jimenez et al., 2014) and mitogen activated kinase kinase (MEK) (Ray et al., 2022) catalyze this process. Shed vesicles contain annexin A1 and annexin A6, though neither is required for shedding (Ray et al., 2022; Ray et al., 2018). Finally, patch repair, the homo/heterotypic fusion of vesicles with the membrane (Cooper and McNeil, 2015), involves patch repair proteins including the muscle-specific dysferlin (Cai et al., 2009; Roostalu and Strahle, 2012), and synaptotagmin 7 (Cooper and McNeil, 2015). The contribution of patch repair to resisting CDCs is not known.

In contrast to CDCs, no repair mechanisms for aerolysin are known. Understanding repair mechanisms to aerolysin is important because it is expected to enable selective targeting of toxins in the future and provide new insights into the generality of repair mechanisms. Repair of cells following aerolysin challenge may take longer than CDCs, potentially due to the small size of the pore (Cooper and McNeil, 2015; Gonzalez et al., 2011; von Hoven et al., 2017).

Furthermore, it is controversial if Ca^2+^ flux is triggered by aerolysin, so it has been questioned if Ca^2+^ dependent membrane repair responses are activated. While early studies suggested aerolysin conducts Ca^2+^ (Krause et al., 1998; Nelson et al., 1999; Tschodrich-Rotter et al., 1996), a recent study reported minimal Ca^2+^ flux following aerolysin challenge (Larpin et al., 2021). However, it was also suggested that Ca^2+^ release by the endoplasmic reticulum (ER) mediates repair (Bittel et al., 2020; Krause et al., 1998; Tschodrich-Rotter et al., 1996).

Alternatively, aerolysin may trigger repair downstream of K^+^ efflux (Gonzalez et al., 2011). K^+^ efflux may trigger repair against the actinoporin sticholysin and CDC listeriolysin O (Cabezas et al., 2017). Thus, it is unclear what repair responses, if any, are activated by aerolysin. Here, we tested the hypothesis that cells use similar repair mechanisms to resist both CDCs and aerolysin. Instead, we found a difference in repair responses. Nucleated cells were 2-4-fold more sensitive to aerolysin than to CDCs. Removal of extracellular Ca^2+^ protected cells from aerolysin, suggesting that Ca^2+^-dependent cell death signals outweigh repair benefits. However, when we removed intracellular Ca^2+^, cells became more sensitive to aerolysin. Live cell imaging showed sustained extracellular Ca^2+^ influx when cells were challenged with aerolysin. This suggested that Ca^2+^-dependent repair was active following aerolysin challenge. We tested four proposed repair pathways: caveolar endocytosis, MEK-dependent shedding, annexin-mediated clogging, and patch repair. We found that caveolar endocytosis was not required for repair against aerolysin or CDCs. MEK-dependent repair was not required to resist aerolysin. Annexin A6 was recruited to the membrane more slowly following aerolysin challenge compared to CDCs. Importantly, cells transfected with dysferlin were more resistant to aerolysin than CDCs. This suggests that cells utilize patch repair as the primary repair mechanism against aerolysin. Overall, these findings suggest that aerolysin and CDCs trigger distinct repair responses, which suggests repair responses are tailored to specific toxin threats.

## Results

### Nucleated cells are more sensitive to Aerolysin than CDCs

To investigate cellular resistance to aerolysin cytotoxicity, and to compare cytotoxicity between aerolysin and CDCs, we normalized toxin activity using human erythrocyte lysis. Using toxin activity in erythrocytes instead of toxin mass allows comparison of relative cellular resistance, and controls for variation in specific activity between toxin preparations. We compared HeLa cell cytotoxicity to SLO, perfringolysin O (PFO), ILY, pro-aerolysin, and trypsin-activated aerolysin (aerolysin) by flow cytometry (Fig 1A, Supplementary Fig S1A-C, S2A). We selected SLO because it is our best-characterized toxin (Haram et al., 2022; Ray et al., 2022; Ray et al., 2018; Romero et al., 2017), and PFO as a second CDC to compare to aerolysin. Finally, we include ILY because it engages the GPI-anchored protein CD59, and aerolysin also engages GPI-anchored proteins. To compare the cytotoxic activity, we performed dose-response curves, calculated toxin specific lysis at each dose, and fit the dose-response curve to a logistic model to determine the toxin dose needed to kill 50% of the cells (LC_50_). To control for impurities in toxin purification, we challenged cells with an equivalent mass of the non-hemolytic aerolysin Y221G (aerolysin*^Y221G^*) (Tsitrin et al., 2002), while we controlled for trypsin activation using trypsin alone (Supplementary Fig S1B). Neither trypsin alone nor aerolysin*^Y221G^* killed cells (Supplementary Fig S1B). Since previous results suggest that aerolysin takes longer to kill cells than CDCs (Cabezas et al., 2017; Gonzalez et al., 2011; Husmann et al., 2009; Ray et al., 2018), we performed a time course to measure aerolysin cytotoxicity. In contrast to SLO, which kills within 5 min post-addition (Keyel et al., 2011; Ray et al., 2018), aerolysin required at least 30 min before robust cytotoxicity was measured (Supplementary Fig S1B, C). Cytotoxicity increased between 30 min and 1 h (Supplementary Fig S1B, C). Since the flow cytometry assay is not suitable for toxin challenge times longer than 1 h, we used an MTT assay to measure viability in independent replicates at 3-24. Cytotoxicity was not different between samples in the first 12 h post-challenge (Supplementary Fig S1D, E). Consistent with previous results (Ray et al., 2018), both PI uptake assay and MTT were comparable to each other (Fig 1A, Supplementary Fig S1, S2A). Based on these results, we chose 30 min as the optimal time for comparison to CDCs. When we compared pro-aerolysin or aerolysin to CDCs, we found that HeLa cells were more sensitive (∼5 fold) to aerolysin, independent of the method of activation (Fig 1A, Supplementary Fig S2A). These data indicate that aerolysin is more toxic than CDCs to HeLa cells.

**Figure 1.**
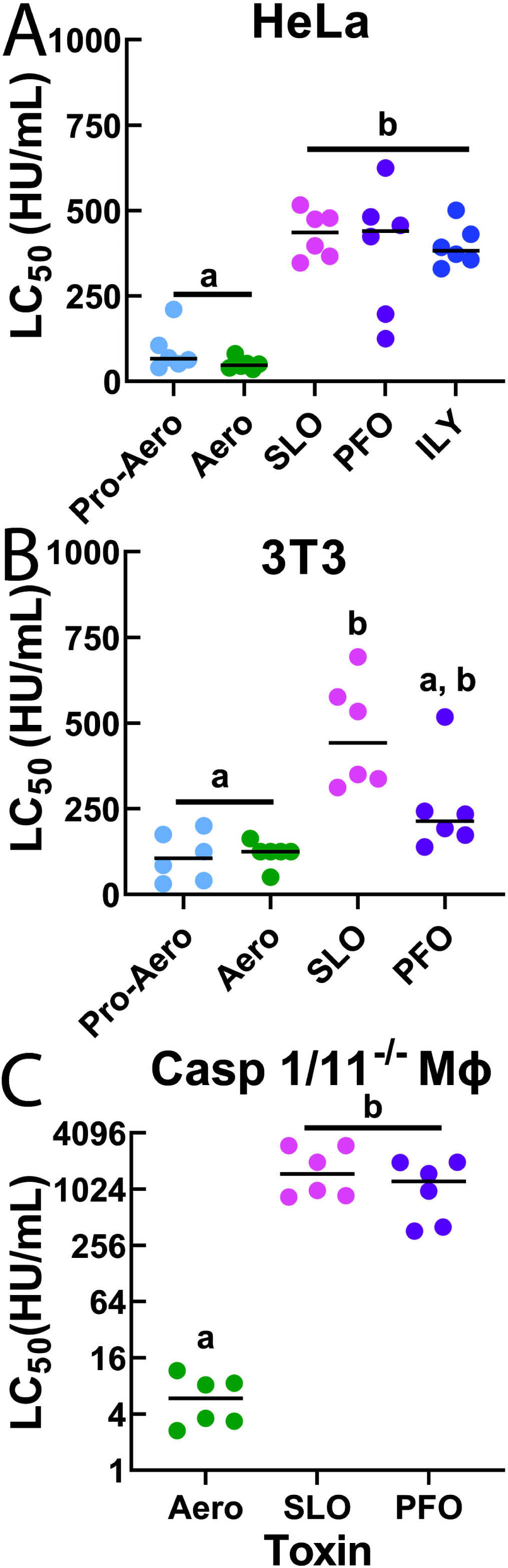
Nucleated cells are more sensitive to aerolysin than CDCs. (A) HeLa, (B) 3T3 or (C) Caspase 1/11^-/-^ bone-marrow derived macrophages (MΦ) were challenged with (A, B) 31-2000 HU/mL pro-aerolysin (Pro-Aero), aerolysin (Aero), streptolysin O (SLO), perfringolysin O (PFO), or (A) intermedilysin (ILY), or (C) 0.5-2000 HU/mL aerolysin, or 15-8000 HU/mL SLO, or PFO in 2 mM CaCl_2_ supplemented RPMI (RC) with 20 μg/ml propidium iodide (PI) for 30 min at 37°C. PI uptake was analyzed by flow cytometry. Specific lysis was determined. The LC_50_ was calculated as described in the methods. Graphs display the median of (A) 8 or (B, C) 6 independent experiments. Data points represent individual experiments. Letters (a-b) denote statistically significant (p < 0.05) groups using repeated-measures analysis of variance (ANOVA) between groups. Thus in 1A, pro-aero and aero are not statistically different from each other (they both are in group a), but are distinct from the CDCs (group b), and vice versa. In 1B, SLO (group b) is statistically significant from pro-aero/aero (group a), but not PFO. Aero is not statistically significant from pro-aero or PFO. PFO is in both groups a and b because it was not statistically significant from either group in the post hoc testing.

To determine if this is a cell-specific phenomenon, we tested two additional cell types: murine fibroblasts (3T3), a model cell line for membrane repair, and Caspase 1/11^−/−^ primary murine bone marrow-derived macrophages (Casp 1/11^−/−^BMDMs). We used Casp 1/11^-/-^ cells to rule out cell death due to inflammasome-dependent Gasdermin D activation (Keyel et al., 2013). The advantage of testing these two cell lines is to give a broader overview of mouse-human differences in sensitivity, and a comparison between immune cells and non-immune cells. When we challenged 3T3 cells with toxins, we found that 3T3 cells were ∼4 to 6 fold more sensitive to pro-aerolysin and aerolysin compared to PFO and SLO, similar to HeLa cells (Fig 1B, Supplementary Fig S2B). In contrast to 3T3 or HeLa cells, Casp 1/11^-/-^ BMDM were ∼100 fold more sensitive to aerolysin compared to SLO and PFO (Fig 1C, Supplementary Fig S2C). The LC_50_ for aerolysin was lower for Casp 1/11^-/-^ BMDM (7.1±1.7 HU/mL) compared to that for HeLa (87.7±8.0 HU/mL) or 3T3 cells (108.7±24.5 HU/mL) (Fig 1, Supplementary Fig S2). This stands in stark contrast to CDCs, where BMDM are ∼5-10 fold more resistant than HeLa cells (Keyel et al., 2013). Based on these data, we conclude that nucleated cells are more sensitive to aerolysin than to CDCs.

### Delayed calcium flux kills aerolysin-challenged cells

The increased sensitivity of aerolysin could be due to reduced repair, or activation of cell death pathways. It is controversial whether cells utilize Ca^2+^- or K^+^-dependent repair responses to resist aerolysin (Cabezas et al., 2017; Gonzalez et al., 2011; Larpin et al., 2021). We tested the impact of ion flux on aerolysin either by chelating extracellular Ca^2+^ with EGTA, or by preventing K^+^ efflux with addition of 150 mM KCl to the extracellular medium. We found no change in the LC_50_ for aerolysin or SLO when K^+^ efflux was blocked (Fig 2A, Supplementary Fig S3A). Consistent with previous results (Ray et al., 2018; Romero et al., 2017), lack of extracellular Ca^2+^ sensitized cells to SLO (Fig 2B, Supplementary Fig S3B). In contrast, removal of extracellular Ca^2+^ protected HeLa cells from pro-aerolysin and aerolysin because the LC_50_ increased ∼10 fold (Fig 2B, Supplementary Fig S3B). To confirm that Ca^2+^ chelation prevented cell death instead of interfering with toxin binding, we measured the binding of Alexa Fluor 647 conjugated pro-aerolysin*^K244C^* (pro-aero*^K244C^*) (Iacovache et al., 2006). Pro-aero*^K244C^* bound equally well to HeLa cells in a dose-dependent manner independently of extracellular Ca^2+^ (Supplementary Fig S3C). Based on these data, we conclude that Ca^2+^ influx, but not K^+^ efflux, is critical for aerolysin-mediated killing.

**Figure 2.**
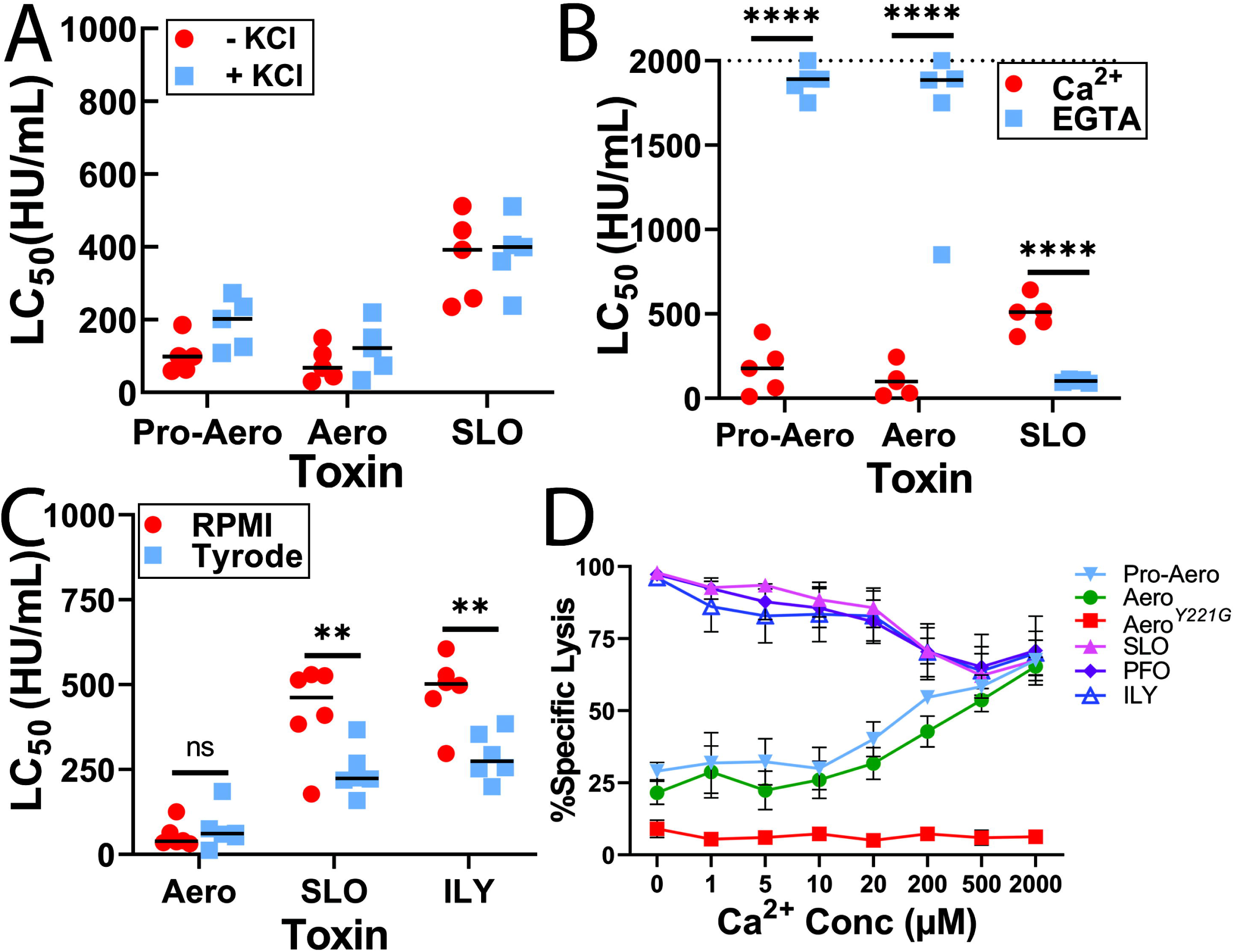
Ca^2+^ influx drives aerolysin cytotoxicity. (A-B) HeLa cells were challenged with 31-2000 HU/mL pro-aerolysin (Pro-Aero), aerolysin (Aero) or SLO for 30 min at 37°C in either RPMI (A) with or without 150 mM KCl, (B) with 2 mM CaCl_2_, or 2 mM EGTA supplemented RPMI. (C) HeLa cells were challenged with 31-2000 HU/mL aerolysin, SLO, or ILY for 30 min at 37°C in RC or Tyrode’s buffer. (D) HeLa cells were challenged with 125 HU/mL pro-aerolysin, aerolysin, 250 HU/mL SLO, PFO or ILY, or mass equivalent to aerolysin of mutant aerolysin^Y221G^ (Aero*^Y221G^*) for 30 min at 37°C in Tyrode’s buffer supplemented with the indicated concentrations of CaCl_2_. PI uptake was analyzed by flow cytometry. The LC_50_ was calculated as described in the methods. (A-C) Data points represent individual experiments. Graphs show the median of (A, B) 5, or (C) 6 independent experiments or (D) the mean ±SEM of 6 independent experiments. Letters (a-c) denote statistically significant (p < 0.05) groups using repeated-measures ANOVA between groups. *** p <0.001, **** p<0.0001.

Since Ca^2+^ is toxic to cells following aerolysin challenge, we determined the amount of extracellular Ca^2+^ necessary for cell death. The medium used in killing assays, RPMI, contains 0.4 mM Ca^2+^, so we used Tyrode’s buffer to control extracellular Ca^2+^. We first compared HeLa cell cytotoxicity between RPMI and Tyrode’s buffer, each supplemented with 2 mM CaCl_2_. Aerolysin was equally potent in both buffers, but SLO and ILY were more potent in Tyrode’s buffer compared to RPMI (Fig 2C, Supplementary Fig S3D). To measure the impact of extracellular Ca^2+^, we measured cytotoxicity at a range of Ca^2+^ concentrations using a toxin dose that killed ∼70% of cells at 2 mM Ca^2+^. Reducing extracellular Ca^2+^ to ∼10 μM protected cells from aerolysin (Fig 2D). Notably, this is consistent with prior results (Iacovache et al., 2016) showing that aerolysin can form pores at low/no Ca^2+^, and suggests EGTA did not interfere with oligomerization or pore formation. In contrast, CDC killing plateaued ∼20 μM (Fig 2D). We conclude that aerolysin cytotoxicity is caused by Ca^2+^ overload.

To better measure Ca^2+^ overload, we labeled cells with fluo-4 and measured calcium influx in HeLa cells challenged with aerolysin or SLO. To analyze Ca^2+^ flux, we categorized cells into three subsets based on their survival time (Ray et al., 2022): cells dying within 5 min of toxin challenge, cells dying within 5-45 min of toxin challenge, and cells surviving throughout the entire 45 min imaging experiment (Fig 3A, Supplementary Fig S3E, Video V1). We chose these categories to stratify cells that were immediately overwhelmed and killed by toxin, cells dying during the course of the imaging, and cells with functioning proximal repair mechanisms. We expect cell death occurring after 45 min to be the result of repair-independent mechanisms, so these processes were not examined. While overall cell death was comparable between SLO and aerolysin, SLO killed more cells within 5 minutes of toxin challenge than aerolysin (Fig 3A), consistent with our findings that aerolysin kills more slowly than SLO (Supplementary Fig S1B, C). In contrast, minimal cell death was observed in cells challenged with aerolysin plus EGTA, non-hemolytic aerolysin*^Y221G^*, or PBS alone (Fig 3A). The maximal Ca^2+^ flux from control cells was lower compared to toxin-challenged cells (Fig 3B, C, Video V2). Notably, the maximal Ca^2+^ flux was close to saturating in our system (fMAX = 1024) with toxin-challenged cells, except for aerolysin-challenged cells that died within 5 min of challenge (Fig 3C, Video V1). When we measured the time to reach the fMAX, all cells that died within 5 minutes rapidly peaked Ca^2+^ (∼4 min) (Fig 3D, E). While surviving cells had a longer time to reach fMAX compared to cells dying between 5 and 45 min, SLO challenged cells reached fMAX faster than aerolysin challenged cells (Fig 3D, F-G). SLO challenged cells showed rapid Ca^2+^ oscillations throughout the imaging period (Video V1). In contrast, aerolysin-challenged cells showed delayed, yet sustained levels of Ca^2+^, with reduced oscillations (Fig 3D, F-G, Video V1). The steady, sustained increase in Ca^2+^ flux caused by aerolysin required pore formation because we did not observe similar effects with cells challenged with PBS, aerolysin with EGTA or aerolysin*^Y221G^* (Fig 3H). We quantitated Ca^2+^ oscillations by determining the realized volatility of the Ca^2+^ flux for each cell. SLO-challenged cells had greater volatility compared to aerolysin-challenged cells (Fig 3I, Supplementary Fig S3F). This analysis showed that the non-hemolytic aerolysin*^Y221G^* also triggered Ca^2+^ oscillations above background, though EGTA reduced Ca^2+^ oscillations to background levels (Fig 3J, Supplementary Fig S3F). These data suggest that pores formed by aerolysin trigger a sustained Ca^2+^ flux that leads to cell death.

**Figure 3.**
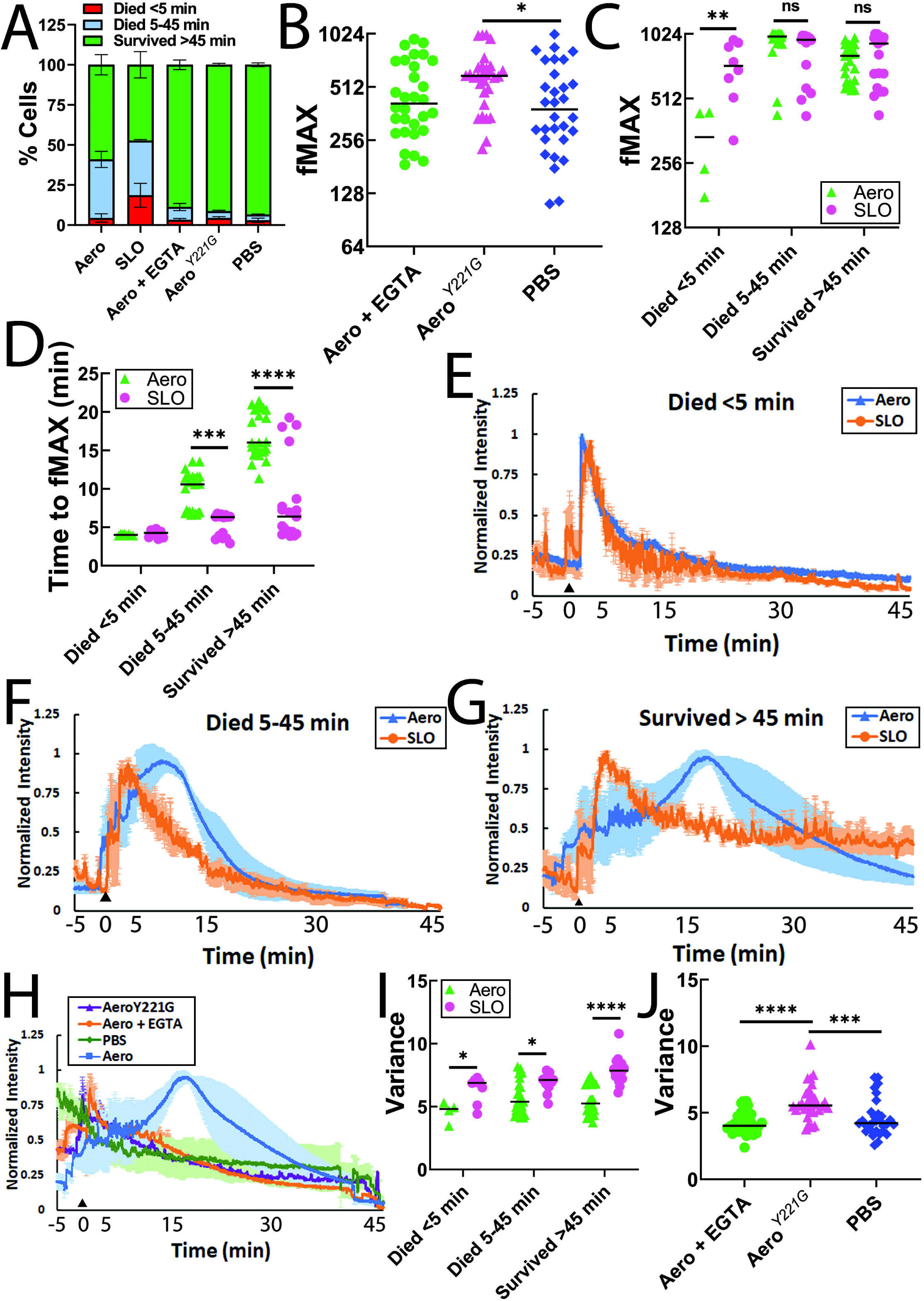
Aerolysin triggers delayed Ca^2+^ influx compared to CDCs. HeLa cells were labeled with Fluo-4 for 30 min in DMEM. The media was replaced with 2 μ TO-PRO3, 25 mM HEPES, pH 7.4, and 2 mM CaCl_2_ supplemented RPMI (imaging buffer). 2 mM EGTA was substituted for CaCl_2_ where indicated. Sublytic toxin (62 HU/mL aerolysin, 250 HU/mL SLO, mass equivalent of aerolysin^Y221G^ (Aero*^Y221G^*)) or PBS was added 5 min after confocal imaging started. Cells were imaged for ∼45 min at 37°C. Ca^2+^ flux and cell death were recorded. Cells were categorized into three subsets by time of cell death: died < 5 min, died between 5 and 45 min, and survived >45 min. (A) The percentage of each subset is shown. (B, C) Maximal Fluo-4 fluorescent intensities (fMAX) are shown for individual cells. (D) The time to reach fMAX is shown. (E-H) Fluo-4 fluorescence intensity was normalized to fMAX for individual cells and averaged. Calcium traces for (E) cells that died < 5 min, (F) cells that died between 5 and 45 min, or (G) cells that survived >45 min after toxin challenge with are shown. (H) Calcium traces for cells challenged with aerolysin with 2 mM EGTA, Aero*^Y221G^* or PBS alone are shown along with the wild type aerolysin trace from (G). Arrowhead indicates toxin addition. (I, J) Ca^2+^ oscillations were quantitated by determining the volatility of the Ca^2+^ flux for each cell. Data points represent individual cells from at least 3 independent experiments. Graphs show the (B-D, I, J) median or (A, E-H) mean ± SEM of at least 3 independent experiments. ns not significant, * p <0.05, ** p <0.01, *** p <0.001, **** p<0.0001.

### Calcium influx does not activate MEK-dependent repair after aerolysin challenge

While extracellular Ca^2+^ is toxic, intracellular Ca^2+^ levels could trigger membrane repair responses. Repair proteins, including annexins, and dysferlin bind Ca^2+^ during repair (Cooper and McNeil, 2015). A prior study suggested intracellular Ca^2+^ can drive repair (Bittel et al., 2020). To test if intracellular Ca^2+^ triggers repair responses to aerolysin, we chelated intracellular Ca^2+^ using BAPTA-AM and challenged cells with aerolysin with or without also chelating extracellular Ca^2+^. Removal of extracellular Ca^2+^ protected cells from aerolysin regardless of intracellular Ca^2+^ chelation (Fig 4A, Supplementary Fig S4A). However, chelation of intracellular Ca^2+^ showed a trend to sensitizing cells to aerolysin (Fig 4A, Supplementary Fig S4A). These data suggest that Ca^2+^ dependent repair pathways could be activated by aerolysin.

**Figure 4.**
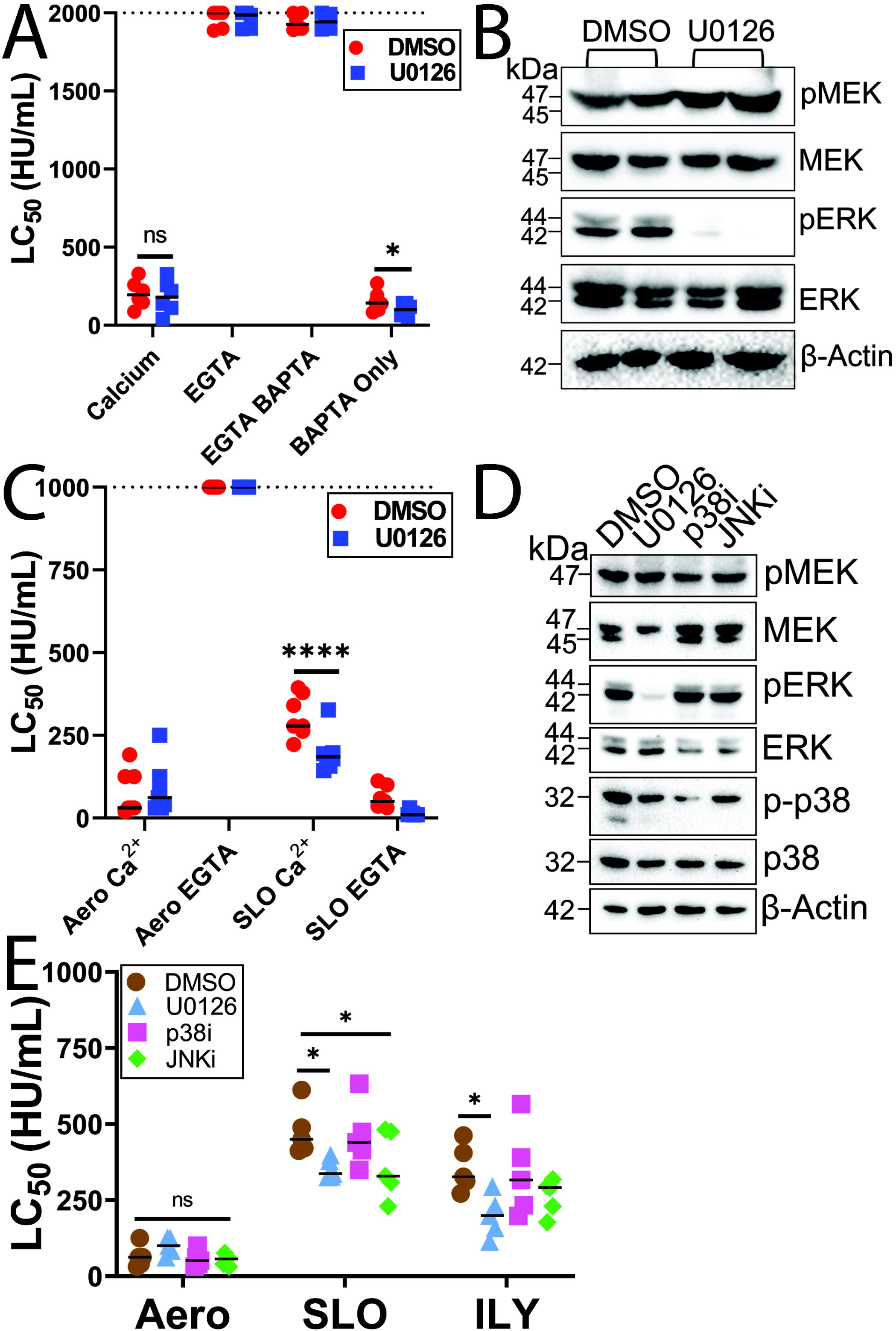
MEK-dependent repair does not contribute to aerolysin resistance. HeLa cells were serum starved for 30 min and pretreated with (A-E) vehicle (DMSO), (A) 100 μM BAPTA-AM, (A-E) 20 μM U0126 (MEK inhibitor), (D, E) 20 μM SB203580 (p38i), or 20 μM SP600125 (JNKi) for 30 min, then either (A, C, E) challenged with toxin or (B, D) lysed for western blot analysis. HeLa cells were challenged in either (A, C) 2 mM EGTA or (A, C, E) 2 mM CaCl_2_ supplemented RPMI with 31-2000 HU/mL of the indicated toxins for 30 min at 37°C. PI uptake was analyzed by flow cytometry. The LC_50_ was calculated as described in the methods. (B, D) Blots were probed with the indicated antibodies and HRP–conjugated secondary antibodies. Graphs show the median of at least (A) 6, (C) 7 or (E) 5 independent experiments. The blots show one representative experiment from (B) 7 or (D) 5 independent experiments. The dotted line indicates the limit of detection. Points on this line had LC_50_ >2000 HU/mL. ns not significant, * p <0.05, **** p<0.0001.

To examine Ca^2+^-dependent repair, we first examined MEK-dependent repair. We recently showed that ∼70% of Ca^2+^-dependent repair of damage caused by CDCs is activated by MEK (Ray et al., 2022). To determine if MEK signaling contributes to survival following aerolysin challenge, we blocked MEK activation with the MEK inhibitor U0126 (Fig 4). U0126 blocked MEK activation (Fig 4B, D), confirming inhibitor activity. We then challenged MEK-inhibited cells with aerolysin or SLO. MEK inhibition did not increase cellular sensitivity to aerolysin (Fig 4, Supplementary Fig S4). As previously observed (Ray et al., 2022), MEK inhibition increased CDC-dependent cell death (Fig 4C, E Supplementary Fig S4B, C). Based on these data, we conclude that MEK-dependent repair is not a major contributor to cell survival from aerolysin.

If MEK is not a major contributor to survival from aerolysin, another could MAP kinase mediate protection. Both p38 and JNK are implicated in long-term viability, protecting cells (Huffman et al., 2004) and recovering ion homeostasis after pro-aerolysin challenge (Gonzalez et al., 2011). We tested if they contribute to membrane repair. Neither p38 nor JNK inhibition altered repair responses to aerolysin or SLO in HeLa cells (Fig 4E, Supplementary Fig S4C). However, JNK inhibition increased sensitivity of HeLa cells to ILY (Fig 4E, Supplementary Fig S4C). Thus, we conclude that MAP kinases are not a major repair mechanism against aerolysin.

### Caveolar endocytosis does not protect cells from aerolysin

We next tested one proposed mechanism of membrane repair: caveolar endocytosis. While we have shown that caveolar endocytosis restores homeostasis instead of mediating repair (Keyel et al., 2011; Romero et al., 2017) following CDC challenge, caveolar endocytosis might mediate aerolysin resistance. To test the role of caveolar endocytosis in aerolysin resistance, we used cells where Cavin-1 (PTRF) was inactivated by CRISPR (Liu and Pilch, 2016). Cavin-1 deficiency prevents caveolae formation (Hill et al., 2008; Liu and Pilch, 2008), and Cavin-1 deficient murine embryonic fibroblasts (MEFs) may have repair defects (Corrotte et al., 2020). We used 3T3-L1 cells instead of MEFs because 3T3-L1 cells can be differentiated to adipocytes, which are richer in caveolae (Scherer et al., 1994; Thorn et al., 2003). If caveolar endocytosis is important for repair, any phenotype is expected to be exacerbated in adipocytes. We differentiated two distinct Cavin-1 deficient 3T3-L1 cell lines (K5 and K3) and one CRISPR-control 3T3-L1 cell line (KN/wild type). To control for any differences due to duration of differentiation time, we compared earlier (Day 9) and later (Day 14) adipocytes. Consistent with previous results (Liu and Pilch, 2016), and similar to HeLa cells, Cavin-1 expression was restricted to 3T3-L1 wild type cells (Fig 5A). To determine the role of caveolar endocytosis in repair in cells with low levels of caveolae, we challenged undifferentiated wild type, K3 or K5 3T3-L1 cells with aerolysin or SLO. Cavin-1 deficiency did not reduce the LC_50_ for either toxin (Fig 5B, Supplementary Fig S5A). These data suggest caveolar endocytosis is not involved in cellular protection or membrane repair against aerolysin or SLO.

**Figure 5.**
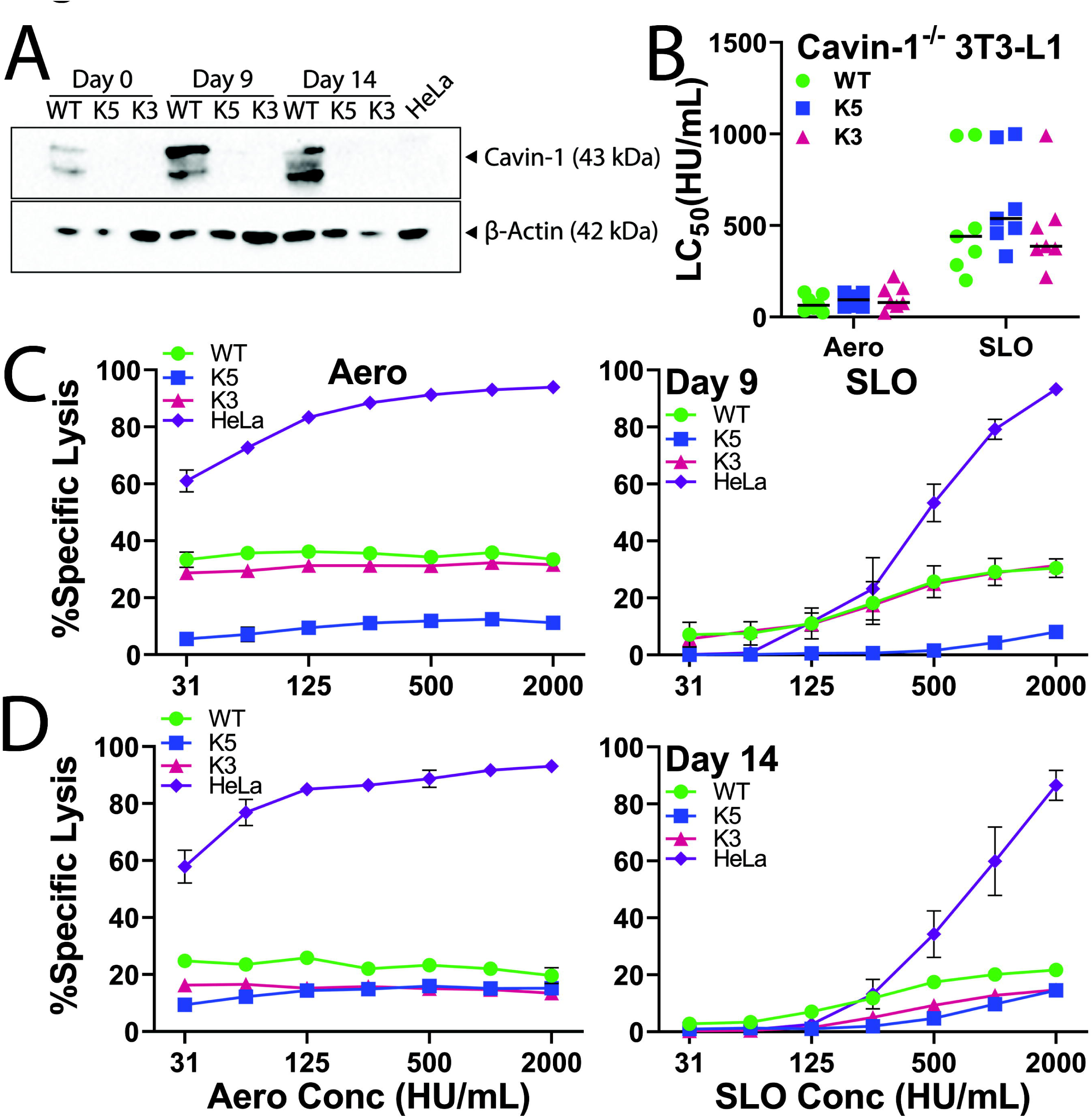
Caveolar endocytosis does not mediate repair responses to aerolysin or CDCs. (A) Wild-type CRISPR control (WT) or two different Cavin-1 (PTRF) CRISPR knockout 3T3-L1 cells (K5 and K3) were undifferentiated or differentiated into adipocytes and collected 0, 9, or 14 days after differentiation. HeLa cells and day 0, 9 or 14 3T3-L1 adipocytes were analyzed by western blot. Portions of the blot were probed with anti-PTRF (anti-Cavin-1) or anti-β-actin antibodies, and HRP-conjugated secondary antibodies. (B-D) WT, K5, or K3 3T3-L1 cells were undifferentiated or differentiated into adipocytes for (C) 9 or (D) 14 days and challenged with 31-2000 HU/mL aerolysin or SLO. PI uptake was analyzed by flow cytometry. The LC_50_ was calculated as described in the methods. The blots show one representative experiment from at least three independent experiments. Graphs show the (B) median of at least 7 or (C, D) mean ± SEM of at least 3 independent experiments.

We next tested cellular resistance under conditions with higher caveolar levels. The number of caveolae increase during adipocyte differentiation (Thorn et al., 2003). We challenged 3T3-L1 cells differentiated into adipocytes for 9 or 14 days with aerolysin or SLO, using HeLa cells as a control. All 3T3-L1 adipocytes were resistant to both SLO and aerolysin, regardless of time point (Fig 5C, D). One Cavin-1^-/-^ line, K5, had increased toxin resistance (Fig 5C) at day 9. However, this did not hold at day 14. The toxins were active because HeLa cells challenged at the same time with the same toxin aliquot remained sensitive (Fig 5C, D). We were unable to calculate the LC_50_ for the adipocytes because the sensitivity was above baseline cell death, but not dose-dependent in the toxin range used. This resistance could be due to an inability of toxins to bind to differentiated adipocytes. To test if adipocytes no longer bind toxins, we challenged HeLa cells or 3T3-L1 adipocytes with fluorescently conjugated pro-aero*^K244C^* or SLO ML (SLO ML Cy5). Both HeLa cells and 3T3-L1 adipocytes bound toxins in a dose dependent manner (Supplementary Fig S5B-E). We interpret these data to indicate that adipocytes are resistant to pore-forming toxins like SLO and aerolysin independently of caveolae. We conclude that caveolar endocytosis does not protect cells from aerolysin or SLO.

### Annexins minimally resist aerolysin

We next examined the ability of annexins to protect cells from aerolysin. To determine the role of annexins in repair against aerolysin, we knocked down three different annexins using RNAi: annexin A1 (A1), annexin A2 (A2), and annexin A6 (A6). We selected these annexins because A2 and A6 are recruited early to plasma membrane damage, while A1 is recruited later, based their calcium sensitivity (Roostalu and Strahle, 2012; Wolfmeier et al., 2015), and we previously characterized their activity during SLO challenge (Ray et al., 2022). The knockdown efficiency was 65-95% (Fig 6A, B). We then challenged these cells with aerolysin and found no significant difference in the sensitivity (Fig 6C, Supplementary Fig S6A). However, knockdown of annexins increased cell sensitivity to SLO, and showed a trend for ILY (Fig 6C, Supplementary Fig S6B-C). These data suggest that annexins are unlikely to contribute to aerolysin resistance to the same degree they contribute to CDC resistance.

**Figure 6.**
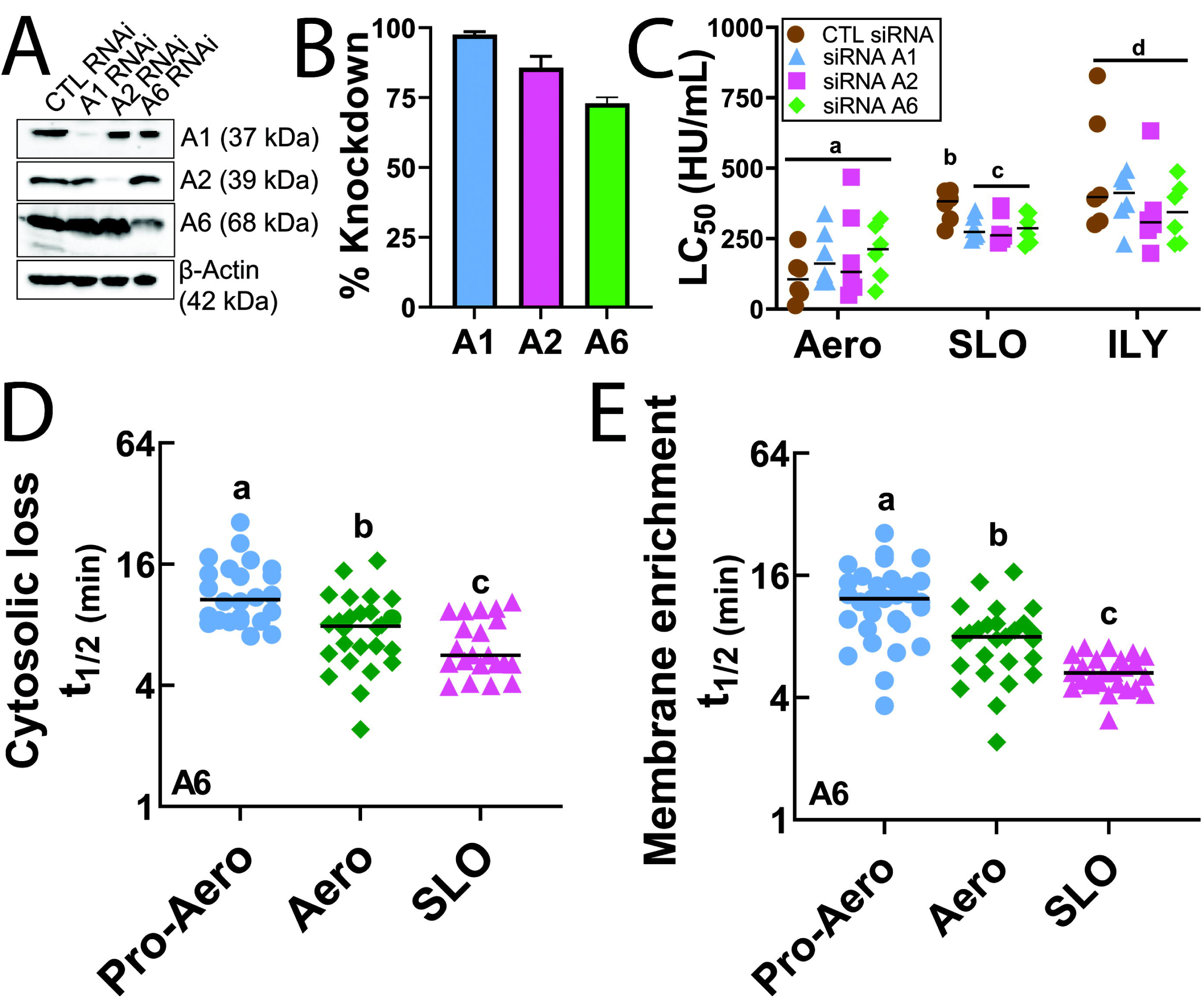
Annexins minimally resist aerolysin. HeLa cells were transfected with control (CTL), A1, A2 or A6 siRNA for 3 days and then (A, B) lysed for western blot analysis, or (C) challenged with 31-2000 HU/mL aerolysin (Aero), SLO or ILY for 30 min at 37°C. PI uptake was analyzed by flow cytometry. (A) Portions of the blot were β-actin followed by HRP-conjugated secondary antibodies. (B) Quantification of the knockdown efficiency compared to control siRNA is shown. (D, E) HeLa cells were transfected with A6-YFP and challenged with sublytic toxin concentrations (62 HU/mL pro-aerolysin or aerolysin, or 250 HU/mL SLO) in imaging buffer. The cells were imaged by confocal microscopy for ∼45 min at 37°C and then lysed with 1% Triton-X. The time to half maximal (t_1/2_) (D) cytosolic depletion or (E) membrane accumulation of A6 is shown. Graphs show the (B) mean ± SEM or (C-E) median. Data points represent (C) individual experiments or (D, E) individual cells from at least three independent experiments. Letters (a-d) denote statistically significant (p < 0.05) groups using repeated-measures ANOVA between groups.

We followed up the annexin RNAi by examining annexins during toxin challenge using high resolution live cell imaging. We measured cytosolic depletion and membrane recruitment of A6-YFP after challenging HeLa cells with a sublytic dose of aerolysin or SLO for 45 min (Fig 6D, E, Video V3). After 45 min, we added 1% Triton as a positive control for membrane translocation and TO-PRO3 uptake. Active toxins, but not aerolysin*^Y221G^*, induced A6 translocation from cytosol to the plasma membrane (Fig 6D, E, Supplementary Fig S7A, B, Video V3). Trypsin activation of aerolysin accelerated A6 cytosolic depletion and membrane translocation (Fig 6D, E, Supplementary Fig S7A, B). A6 cytosolic depletion took 10 min for aerolysin, or 15 min for pro-aerolysin compared to ∼6 min for SLO (Fig 6D). The faster cytosolic loss of A6 for SLO was paralleled by a faster membrane recruitment of A6 (∼5 min) (Fig 6E, Supplementary Fig S7A, B). We found a similar trend of membrane enrichment of A6 for aerolysin (∼10 min) or pro-aerolysin (∼15 min) (Fig 6E). Since annexin depletion does not measure viability or permeability, we analyzed TO-PRO3 uptake. We found that all toxins induced A6-YFP translocation without TO-PRO3 accumulation (Supplementary Fig S7C, Video V3).

### Microvesicle shedding is reduced during aerolysin challenge

We next analyzed microvesicle shedding by toxin challenged cells. As done in previous studies (Babiychuk et al., 2009; Ray et al., 2022; Ray et al., 2018), we used A6 as a marker for microvesicle shedding. Pro-aerolysin and aerolysin induced ∼2-fold lower shedding of A6 enriched microvesicles compared to SLO (Fig 7A). Trypsin treatment of aerolysin did not change the rate of microvesicle shedding (Fig 7A). Aerolysin-triggered shedding was increased compared to shedding by non-hemolytic, oligomeric aerolysin*^Y221G^*, which was minimal (Fig 7A). These data are consistent with the inability of aerolysin to activate MEK-dependent repair (Fig 4). We interpret these data to suggest that aerolysin may activate the same MEK-independent shedding observed with CDCs (Ray et al., 2022). Thus, aerolysin does not trigger microvesicle shedding to the same extent as CDCs.

**Figure 7.**
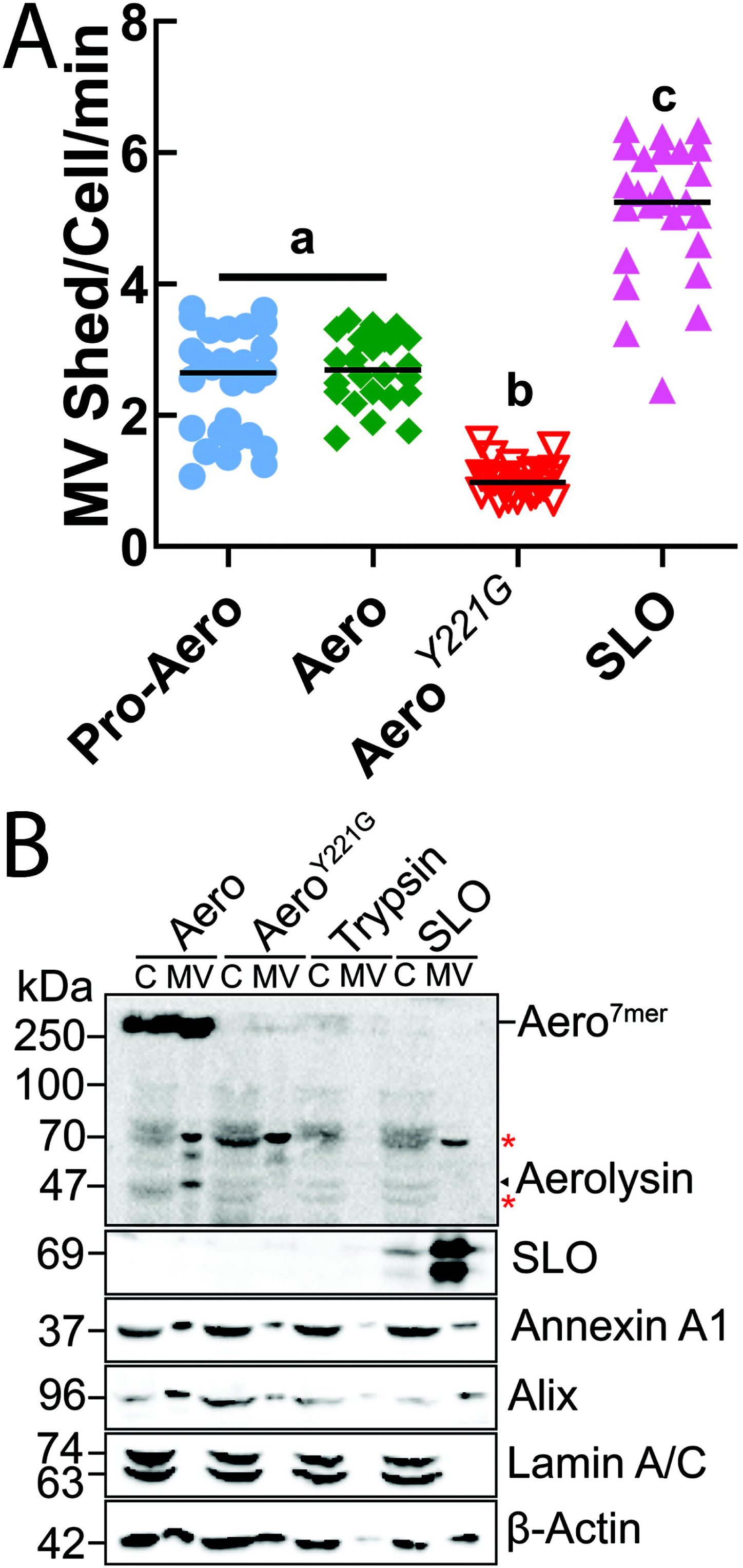
Aerolysin triggers limited shedding. (A) A6-YFP transfected HeLa cells were challenged with a sublytic dose (62 HU/mL) of pro-aerolysin (Pro-Aero) or aerolysin (Aero), a mass equivalent of Aero*^Y221G^* or sublytic dose (250 HU/mL) of SLO in imaging buffer. The cells were imaged by confocal microscopy for ∼45 min at 37°C and then lysed with 1% Triton-X. Shedding of annexin A6^+^ microvesicles was manually counted and expressed as microvesicles shed/cell/min. (B) HeLa cells were challenged for 15 min at 37°C with sublytic doses of aerolysin, SLO, mass equivalent of Aero*^Y221G^* or sufficient trypsin to activate pro-aerolysin. Cells were pelleted at 2000×g for 5 min to yield cell pellet (C). Cell supernatants were spun at 100,000×g for 40 min at 4°C and MV pellet (MV) collected. Fractions were analyzed by western blot using anti-aerolysin, 6D11 anti-SLO, CPTC-A1–3 anti-annexin A1, anti-Alix, MANLAC-4A7 anti-Lamin A/C or AC-15 anti-β-Actin. Graph shows the mean ± SEM. Data points represent individual cells from at least three independent experiments. The blots show one representative experiment from three independent experiments. Letters (a-c) denote statistically significant (p < 0.05) groups using repeated-measures ANOVA between groups. * indicates non-specific bands.

We next examined microvesicle shedding by ultracentrifugation and western blot. We isolated microvesicles shed by non-transfected HeLa cells following toxin challenge by ultracentrifugation. To control for spontaneously released vesicles, we included an equivalent amount of trypsin used to activate aerolysin. We detected aerolysin using an affinity-purified custom rabbit polyclonal antibody raised against aerolysin*^Y221G^*. We validated the antibody specificity against both aerolysin and aerolysin*^Y221G^* by western blotting, ELISA, and immunofluorescence (Supplemental Fig S8). We found that aerolysin and SLO were both shed on MV (Fig 7B). Aerolysin*^Y221G^* localized at low levels on MV (Fig 7B). Notably, SLO was shed to a greater degree than aerolysin, consistent with our A6 shedding results (Fig 7A). Along with active toxins, we found that the membrane repair proteins A1 and Alix were shed in microvesicles (Fig 7B), consistent with the previous results (Jimenez et al., 2014; Romero et al., 2017). To control for cellular debris from lysed cells in our shed microvesicles, we probed for nuclear protein Lamin A/C. We found that Lamin A/C localized to cell pellet only (Fig 7B), indicating that the microvesicle fraction did not contain detectable cellular debris. Trypsin alone did not induce shedding of any repair proteins (Fig 7B). These findings confirm that cells shed microvesicles following aerolysin challenge, but at a reduced extent compared to CDCs. Based on these data, we conclude that microvesicle shedding is not the primary mechanism used to resist aerolysin pores.

### Patch repair protect cells from aerolysin

Finally, we tested the role of patch repair in resisting aerolysin. Due to the redundancies in patch repair, we used a gain-of-function approach. We used exogenous expression of the well-characterized muscle patch repair protein dysferlin in HeLa cells to measure increases in patch repair. We transfected HeLa cells with GFP or GFP-dysferlin, and challenged cells with toxins. GFP-transfection did not significantly increase cellular resistance to aerolysin, SLO or ILY (Fig 8A, B Supplementary Fig S9). Dysferlin increased SLO resistance beyond GFP alone, but was not effective in protecting cell integrity against another CDC, ILY (Fig 8A). Dysferlin increased cellular resistance to aerolysin 4-18 fold (Fig 8A, B Supplementary Fig S9). To confirm the Ca^2+^ dependence of dysferlin-mediated protection, we challenged transfected cells with or without extracellular Ca^2+^. Dysferlin-mediated repair was dependent on Ca^2+^ (Fig 8B). Thus, dysferlin expression protects cells against aerolysin.

**Figure 8.**
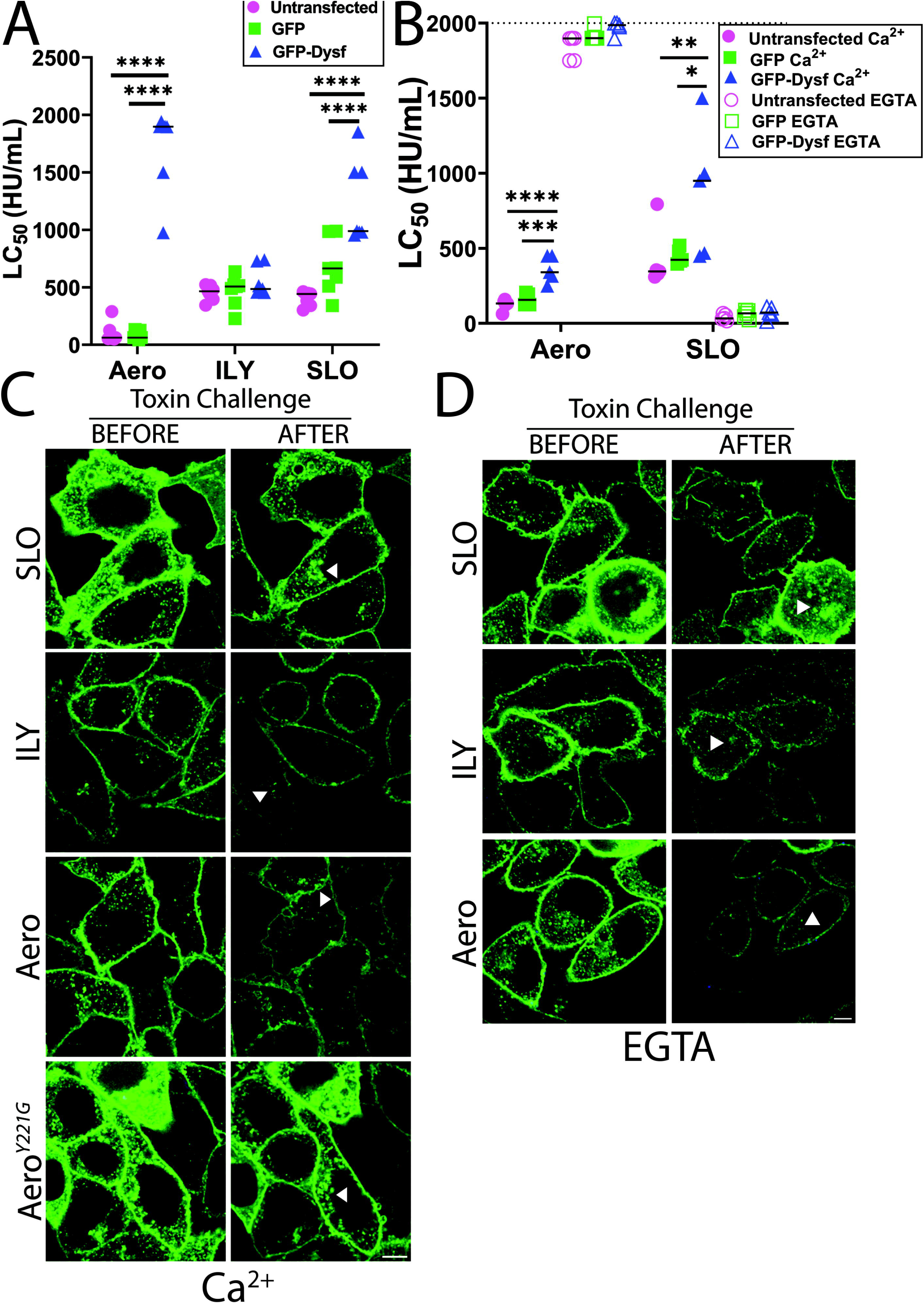
Patch repair protects cells from aerolysin. HeLa cells were either untransfected or transfected with GFP or GFP Dysferlin (GFP-Dysf) for 48 h and challenged with aerolysin (Aero), SLO or ILY at 31-2000 HU/mL for 30 min at 37°C either in (A, B) 2 mM CaCl_2_ or (B) 2 mM EGTA supplemented RPMI. PI uptake was analyzed by flow cytometry. The LC_50_ was calculated as described in the methods. (C, D) GFP-Dysf (green) transfected HeLa cells were challenged with sublytic toxin doses (250 HU/mL SLO or ILY, 62 HU/mL aerolysin or mass equivalent to aerolysin of mutant Aero*^Y221G^*) in imaging buffer either supplemented with (C) 2 mM CaCl_2_ or (D) 2 mM EGTA, imaged for ∼45 min at 37°C, and then lysed with 1% Triton-X. Depletion of GFP-Dysf containing vesicles from cytosol are shown by arrowheads. Graphs show the median of (A) 7, or (B) 5 independent experiments. The dotted line indicates the limit of detection. Points on this line had LC_50_ >2000 HU/mL. Micrographs show representative images from three independent experiments. * p<0.05, ** p <0.01, *** p <0.001, **** p<0.0001. Scale bar = 5 μm.

While dysferlin is associated with patch repair, it might not mediate resistance via patch repair. To confirm that dysferlin mediated patch repair, we measured GFP and GFP-dysferlin dynamics during toxin challenge by high resolution imaging. GFP was cytosolic, and active toxins depleted GFP from the cell over time (Video V4). GFP-dysferlin localized to the plasma membrane and vesicles independently of extracellular Ca^2+^ (Fig 8C D, Video V5). When challenged with sublytic aerolysin dose in presence of Ca^2+^, we found that dysferlin-enriched vesicles moved to the plasma membrane from cytosol and protected the integrity of plasma membrane by patching and blebbing (Fig 8C, Video V5). Non-hemolytic oligomeric aerolysin*^Y221G^* did not trigger dysferlin-mediated patching (Fig 8C, Video V5). To study the Ca^2+^ dependency of dysferlin-positive vesicle fusion with the plasma membrane, we removed extracellular Ca^2+^ with 2 mM EGTA and challenged with sublytic toxin doses. We found less depletion of dysferlin-positive vesicles from cytosol (Fig 8D, Video V6), and cells were more resistant to aerolysin, confirming our previous results (Fig 2B, 4A, C). When challenged with sublytic SLO or ILY in presence of EGTA, cells showed larger bleb formation and less dysferlin-positive vesicle transport to the plasma membrane (Fig 8D, Video V6). However, at later stages dysferlin-positive vesicles moved from the cytosol to plasma membrane, suggesting that intracellular Ca^2+^ could activate patch repair as a last resort. Based on these data, we conclude that patch repair is the primary mechanism used to resist aerolysin pores.

## Discussion

In this study, we demonstrate that patch repair is the primary repair response used by cells to protect themselves against the small pore forming toxin aerolysin. Aerolysin is more toxic to nucleated cells compared to CDCs. The increased susceptibility to aerolysin depends on Ca^2+^ toxicity, and the limited activation of protective repair pathways. Caveolar endocytosis did not protect cells against any toxin we tested. However, intracellular Ca^2+^ activated delayed annexin recruitment, limited shedding, and patch repair. Of these repair mechanisms, patch repair was the primary repair response, which contrasts with CDCs. This suggests that toxins trigger distinct membrane repair pathways downstream of Ca^2+^ flux. Overall, our results provide clarity into repair responses activated by aerolysin.

We found that aerolysin was more potent against nucleated mammalian cells compared to CDCs. In order to compare toxins with distinct sizes, subunits in the complete pore, and relative killing capacity, we normalize cytotoxicity using hemolytic activity. Since erythrocytes are not thought to have any specific membrane repair mechanisms, they provide a baseline for toxin activity. When normalized to equal hemolytic activity, we found that cells are highly sensitive to aerolysin compared to CDCs. In contrast to SLO, but similar to PFO, we found that sensitivity increased over time (Ray et al., 2018). Our results are consistent with previous results suggesting that aerolysin is a potent toxin (Gonzalez et al., 2011; Larpin et al., 2021). Thus, these findings suggest that aerolysin triggers minimal repair compared to CDCs.

While limited repair is one possible interpretation of the results, another possibility is that aerolysin triggers a death pathway that overcomes any repair pathways. We investigated this possibility by controlling ion flux in cells challenged with aerolysin or CDCs. In contrast to prior results suggesting that K^+^ depletion was important during aerolysin challenge (Cabezas et al., 2017), we found no impact on cell survival at 30 min when blocking K^+^ depletion. However, we measured acute responses in HeLa cells, whereas other studies looked at longer time (1-6 h) in HT29 (Gonzalez et al., 2011) or BHK (Cabezas et al., 2017) cells. While prior studies did not report hemolytic activity of their toxin preparations, we used a higher mass of toxin, potentially due to differences in purification. In our system, we found that Ca^2+^ influx was essential for aerolysin cytotoxicity.

Compared to CDCs, aerolysin triggered a delayed and sustained Ca^2+^ flux characterized by few oscillations, as measured by volatility. We interpret the sustained and limited volatility influx as the activation of cellular Ca^2+^ conducting protein(s) and/or lipids. In contrast, we interpret the CDC-driven volatility as the cell attempting to control intracellular Ca^2+^ concentrations, possibly due to membrane resealing, or possibly attempting to reverse activation of cellular Ca^2+^ conducting protein(s) and/or lipids. These results are consistent with previous findings (Krause et al., 1998) supporting aerolysin-dependent Ca^2+^ flux. However, our results diverge from a previous study (Larpin et al., 2021) that used less toxin and did not detect Ca^2+^ flux in THP.1 or U937 cells. One possibility is that THP.1 and U937 cells are unable to activate the cellular Ca^2+^ conducting protein(s) and/or lipids. The activation of a Ca^2+^ death pathway could also obscure activation of repair mechanisms against aerolysin. Overall, activation of a cell type specific Ca^2+^ death pathway that obscures any repair could reconcile conflicting interpretations (Krause et al., 1998; Larpin et al., 2021) of the impact of Ca^2+^ influx on aerolysin. However, it remains to be determined how Ca^2+^ mediates toxicity in aerolysin challenged cells.

We determined which repair mechanisms triggered by CDCs were activated in response to aerolysin. Caveolar endocytosis has been proposed as a repair mechanism (Corrotte et al., 2020), but we found caveolar endocytosis did not promote repair against either aerolysin or CDCs. However, we tested pre-adipocyte and adipocyte cell lines instead of MEFs. Cavin-1 could be required for the membrane homeostasis of cholesterol or other key repair lipids/proteins in MEFs. Notably, intact toxin pores perforating the membrane have been observed on shed blebs (Keyel et al., 2011) but not in the membrane of internal organelles. Our findings here are consistent with our prior finding (Romero et al., 2017) that caveolar endocytosis does not contribute to repair, but may restore cellular homeostasis by removing inactive toxin and toxin laden blebs after repair is complete.

Aerolysin, unlike CDCs, was not resisted by MEK dependent repair. Since MEK dependent repair accounted for 70% of the repair against CDCs (Ray et al., 2022), failure of this mechanism to restrict damage could account for the increased cytotoxicity of aerolysin. We also tested p38 and JNK activation because they were implicated in restoring ion-homeostasis and long-term viability following aerolysin challenge (Gonzalez et al., 2011; Huffman et al., 2004). In contrast to previous results (Gonzalez et al., 2011; Huffman et al., 2004), we did not find changes in repair responses after blocking p38 or JNK MAPK pathways. This difference could be due to the differences in toxin preparation, duration of toxin challenge, or the difference in time frame we used (30-60 min), compared to previous studies (6+ hours/days). Overall, this suggests that p38 and JNK do not contribute to proximal membrane repair events.

We next compared the ability of three annexins (A1, A2 and A6) to promote repair following CDC or aerolysin challenge. Deletion of A1, A2 or A6 did not increase cellular sensitivity to aerolysin, but did sensitize cells to SLO and ILY. This suggests that annexins either serve redundant functions in aerolysin resistance, or A1, A2 and A6 are minimally involved. We propose they are minimally involved because they also showed delayed translocation. The delayed translocation of annexins is consistent with the slower increase in Ca^2+^ that we observed. Delayed, steady Ca^2+^ influx could account for the failure of annexins to contribute to protecting cells from aerolysin. It remains possible that other annexins resist aerolysin. It remains unknown how previously described interactions of annexins with other repair proteins, including dysferlin, MG53, and ESCRT III (Bittel et al., 2020; Demonbreun et al., 2016; Lennon et al., 2003; Roostalu and Strahle, 2012; Sonder et al., 2019), impact repair.

While annexins are associated with clogging, they also mark cellular shedding from the membrane. Shedding of A6^+^ microvesicles was reduced by approximately half when challenged with aerolysin compared to SLO. Since we observed a 2-fold reduction in shedding when MEK was inhibited (Ray et al., 2022), we interpret these results to indicate that aerolysin activates the MEK-independent shedding also observed with CDCs. Ultracentrifugation confirmed that shedding occurs, despite being reduced. Whether the MEK-independent repair is lipid-mediated (Keyel et al., 2011) or ESCRT-mediated (Jimenez et al., 2014), remains to be determined. However, a low contribution of repair by MEK and annexins is consistent with the enhanced cytotoxicity of aerolysin.

In contrast to other repair pathways, we found that enhancing patch repair made cells resistant to aerolysin. Overexpression of the muscle patch repair protein dysferlin protected cells from aerolysin, to a lesser extent SLO, and not to ILY. Interestingly, both ILY and aerolysin target GPI-anchored proteins, though ILY is restricted to human CD59 (Abrami et al., 1998b; Giddings et al., 2004). We did not rule out a direct effect of dysferlin on toxic Ca^2+^ levels. Dysferlin could reduce Ca^2+^ toxicity in aerolysin challenged cells either by sequestering excess cytoplasmic Ca^2+^ with its seven Ca^2+^-binding C2 domains, as reported in t-tubules (Kerr et al., 2013) or by blocking the activation of the Ca^2+^-dependent death effector protein(s) and/or lipids. However, two lines of evidence suggest that patch repair is the mechanism of action in our system. First, survival did not correlate with the extent of dysferlin overexpression, which would be expected if dysferlin acted as a Ca^2+^ sink. For example, very bright dysferlin positive cells were not more resistant than dim, dysferlin positive cells. Second, we observed depletion of dysferlin-positive vesicles as they fused with the plasma membrane during toxin insult. Thus, we conclude that patch repair is the primary resistance mechanism against aerolysin.

One new horizon opened by our work is determining how dysferlin-enhanced patch repair prevents aerolysin induced death. While vertex fusion (McNeil and Kirchhausen, 2005) displacing aerolysin pores from the plasma membrane is one possibility, another possibility is that endolysosomal proteins and/or lipids delivered to the membrane clog or remove the aerolysin pore. Alternatively, dysferlin and/or endolysosomal components could interfere with the Ca^2+^-dependent death pathway via an unknown mechanism.

Our work had limitations. While we showed Ca^2+^ is toxic to cells after aerolysin challenge and removing Ca^2+^ with EGTA protected cells, the signaling intermediate between Ca^2+^ influx and the death effector remains unknown. We used exogenous expression of dysferlin to study patch repair, but did not knock out dysferlin. Future investigation is needed in muscle cells where dysferlin is deleted to compare resistance. Finally, we focused on three key repair responses, but did not examine ESCRT-mediated repair, sphingomyelin/ceramide, other annexins or other repair proteins. These factors could also contribute to repair against aerolysin. Similarly, while this work builds a foundation on which to investigate interactions between repair pathways, we did not pursue potential overlap or redundancy of pathways.

Overall, the key contribution of this study is the comparison of multiple repair responses, and determining which is used by cells against the small PFT aerolysin. While MEK-independent microvesicle shedding was observed, neither MEK nor annexins appear to drive major cellular resistance. Both of these responses were reduced compared to responses against CDCs. Since CDCs are less potent than aerolysin against nucleated cells, we propose aerolysin evades the key Ca^2+^-dependent repair mechanisms shedding and annexin recruitment. However, patch repair is activated as a last resort when other repair mechanisms fail to reseal the damage.

## Materials and Methods

### Reagents

All reagents were from Thermo Fisher Scientific (Waltham, MA, USA) unless otherwise noted. The MEK1/2 inhibitor U0126 was from Cell Signaling Technology (Danvers, MA, USA) (Cat# 9903S) or Tocris, (Minneapolis, MN, USA) (Cat# 1144). The p38 MAPK inhibitor SB203580 (Cat # AG-CR1-0030-M005) was from AdipoGen Life Sciences (San Diego, CA, USA). The MAPK9/JNK1/2 inhibitor SP600125 (Cat # S1460) was from Selleckchem (Houston, TX, USA). siRNAs for annexin A1 and A2 were designed and ordered from Cytiva (Marlborough, MA, USA). Negative control siRNA (Cat # 462001) and annexin A6 siRNA were from Ambion (Cat# AM16708). BAPTA-AM (Cat # 15551) was from Cayman Chemical (Ann Arbor, MI, USA). Propidium iodide (PI) was from Sigma-Aldrich (St. Louis, MO, USA) (Cat# P4170-100MG) or Biotium (Fremont, CA, USA) (Cat # 40016). DAPI (Cat# D9542), Dexamethasone (Cat# D2915-100MG) and insulin (Cat# I6634-50MG) were from Sigma-Aldrich. 3-Isobuty-1-methylxanthine (IBMX) (Cat# 195262) was from MP Biomedicals (Solon, OH, USA). MTT reagent (Cat# 2809-1G) was from BioVision (Milpitas, CA, USA).

### Commercial Antibodies

The anti-annexin A1 monoclonal antibody (mAb) (Clone: CPTC-ANXA1-3-s) was deposited to the Developmental Studies Hybridoma Bank (DSHB) by Clinical Proteomics Technologies for Cancer (DSHB Hybridoma Product CPTC-ANXA1-3) and anti-Lamin A/C (MANLAC-4A7-s) was deposited to the DSHB by GE Morris (DSHB Hybridoma Product MANLAC1(4A7)). Both were obtained from the DSHB created by the NICHD of the NIH and maintained at the University of Iowa, Department of Biology (Iowa City, IA, USA). Anti-β-actin AC-15 mAb (Cat# GTX26276) was from GeneTex (Irvine, CA, USA). Anti-SLO antibody 6D11 mAb (catalog: NBP1–05126) was from Novus Biologicals (Littleton, CO, USA). Anti-Alix 3A9 mAb was from BioLegend (San Diego, CA, USA). Anti-PTRF rabbit polyclonal Ab (Cat# ab48824) was from Abcam (Cambridge, MA, USA). Anti-MEK (9122L), anti–phospho-MEK-[Ser^217^/Ser^221^] (9121S), anti-ERK p44/42 (9102S), anti-phospho-ERK[Thr^202^/Tyr^404^] (9101S), anti-p38 (9212S), and anti–phospho-p38 (94511S), rabbit polyclonal antibodies were from Cell Signaling Technologies (Danvers, MA, USA). Anti-mouse (711-035-151) and anti-rabbit (711-035-152) HRP-conjugated antibodies were from Jackson Immunoresearch (West Grove, PA, USA).

### Aerolysin antibody

Non-hemolytic, oligomeric aerolysin Y221G (Aero*^Y221G^*) was purified and concentrated to 5 mg/mL at 95% purity. Two New Zealand White rabbits were immunized and bled by Pacific Immunology (Ramona, Ca, USA), as overseen by their Institutional Animal Care and Use Committee. After bleeding to collect pre-immune serum, rabbits were immunized once in complete Freund’s adjuvant, a second time on day 21 in incomplete Freund’s adjuvant, a third time on day 42 in incomplete Freund’s adjuvant, and a fourth time on day 70 in incomplete Freund’s adjuvant. Antisera was collected on days 49, 63, 77, 91, with final exsanguination bleeds on days 98 and 101. The antisera was affinity purified against pro-aerolysin. For affinity purification, pro-aerolysin was dialyzed with coupling buffer overnight at 4°C. Then, pro-aerolysin was coupled to cyanogen bromide activated sepharose beads for 2 h at room temperature (RT) and washed with coupling buffer. Remaining sites on the sepharose were quenched using 0.1 M Tris-HCl (pH 8) for 2 h at RT. Coupled beads were then washed alternatively with acid wash (0.1 M sodium acetate, 0.5 M sodium chloride, pH 4) and alkali wash (0.1 M Tris, 0.5 M sodium chloride, pH 8) buffers four times. Anti-aerolysin antiserum was then incubated with beads for 2 h at RT and the mixture loaded on an empty column. The flowthrough was collected and saved as depleted serum. Beads were washed with PBS followed by 150 mM NaCl, and A_280_ was measured until it reached to <0.01. Once the A_280_ was <0.01, high-affinity antibodies were eluted with 0.1 M glycine elution buffer (0.1 M glycine, pH 2.5 with HCl) and collected into 1 mL fractions with 50 µL 1 M Tris (pH 8) until A_280_ was low. The column was washed with 0.1 M Tris. Next, antibodies were again eluted into 1 M Tris using 0.1 M triethylamine (TEA) elution buffer (0.1 M TEA, pH 11.5). For each fraction, the antibody concentration was calculated using 1.4 A_280_ = 1 mg/mL antibody. Relevant elutions were pooled together and final antibody concentration was calculated.

### Plasmids

The pET22b plasmids encoding His-tagged aerolysin and aerolysin*^Y221G^* were kind gifts from Gisou van der Goot (École Polytechnique Fédérale de Lausanne, Canton of Vaud, Switzerland) (Tsitrin et al., 2002). The wild type aerolysin contained Q254E, R260A, R449A and E450Q mutations and the C-terminal KSASA was replaced with NVSLSVTPAANQLE HHHHHH compared to sequences in GenBank (i.e. M16495.1). The K244C mutation was introduced into wild type aerolysin by Quikchange mutagenesis. Cys-less, His-tagged, codon-optimized SLO C530A and “monomer-locked” SLO G398V/G399V (SLO ML) were previously described (Ray et al., 2022). Cys-less His-tagged PFO in pET22 (Shepard et al., 1998) and Cys-less His-tagged ILY in pTrcHisA (Giddings et al., 2004) were generous gifts from Rodney Tweten (University of Oklahoma Health Sciences Center, Oklahoma City, OK, USA). Human Annexin A6 fused to YFP was a generous gift from Annette Draeger (University of Bern, Bern, Switzerland) (Babiychuk et al., 2009). GFP-dysferlin was cloned into pcDNA4.

### Mice

All experimental mice were housed and maintained according to Texas Tech University Institutional Animal Care and Use Committee (TTU IACUC) standards, adhering to the Guide for the Care and Use of Laboratory Animals (8th edition, NRC 2011). TTU IACUC approved mouse use. Casp1/11^-/-^ mice on the C57BL/6 background were purchased from the Jackson Laboratory (Bar Harbor, ME, USA) (stock #016621). Mice of both sexes aged 6–15 weeks were used to prepare BMDM. Sample size was determined as the minimum number of mice needed to provide enough bone marrow for experiments. Consequently, no randomization or blinding was needed. Mice were euthanized by asphyxiation through the controlled flow of pure CO_2_ followed by cervical dislocation.

### Cell culture

All cell lines were maintained at 37°C, 5% CO_2_. HeLa cells (ATCC (Manassas, VA, USA) CCL-2) were cultured in DMEM (Corning, Corning, NY, USA) supplemented with 10% Equafetal serum blend (Atlas Biologicals, Fort Collins, CO, USA) and 1x L-glutamine (D10 medium). 3T3 and 3T3-L1 cells (ATCC, CL-173) were cultured in D10 medium supplemented with 1 mM sodium pyruvate (Corning) and 1x non-essential amino acids (GE Healthcare, Pittsburgh, PA, USA). 3T3-L1 cell lines with PTRF/Cavin-1 inactivated by CRISPR (K5 and K3) and the CRISPR control 3T3-L1 cell line (KN/ wild type) were generous gifts from Libin Liu (Liu and Pilch, 2016). 3T3-L1 cells were differentiated to adipocytes for 3 days in Adipocyte Differentiation media (D10 medium supplemented with 1 µM dexamethasone, 0.5 mM 3-Isobutyl-1-methylxanthine, 10 µg/ml insulin) and maintained for the next 10 days in Adipocyte Maintenance media (D10 medium supplemented with 10 µg/ml Insulin) by changing media every other day. We used adipocytes after 9 or 14 days of differentiation for assays. Casp1/11^-/-^ BMDM were isolated from bone marrow and cultured as previously described (Ray et al., 2022; Romero et al., 2017). BMDM were differentiated for 7-21 days in DMEM supplemented with 30% L929 cell supernatants, 20% fetal calf serum (VWR Seradigm, Radnor, PA, USA), 1 mM sodium pyruvate and 1x L-glutamine.

### Recombinant toxins

Toxins used in assays were induced in *E. coli* BL21 GOLD cells and purified as previously described (Keyel et al., 2012a; Ray et al., 2022; Ray et al., 2018; Romero et al., 2017). Briefly, toxins were induced with 0.2% arabinose (SLO, SLO ML), or 0.2 mM IPTG (PFO, ILY, or pro-aerolysin and pro-aerolysin*^Y221G^*) for 3 h at room temperature, frozen, lysed, and then purified using Nickel-NTA resin. For antibody production, or to determine the impact of purity on toxin activity, pro-aerolysin or pro-aerolysin*^Y221G^* were further purified by FPLC Q anion exchange chromatography (Supplementary Fig S10A). The protein concentration was determined by Bradford Assay (Table 1). Purity was assessed by SDS-PAGE. The hemolytic activity of each toxin was determined as previously described (Keyel et al., 2012a; Ray et al., 2022; Ray et al., 2018; Romero et al., 2017) using human red blood cells (Zen-Bio, Research Triangle Park, NC, USA) (Table 1). One hemolytic unit (HU) is defined as the quantity of toxin required to lyse 50% of a 2% human red blood cell solution in 30 min at 37 °C in 2 mM CaCl_2_, 10 mM HEPES, pH 7.4 and 0.3% BSA in PBS (Keyel et al., 2012a; Romero et al., 2017). In effect, 1 HU is the LC_50_ of the toxin against human red blood cells. To determine the optimal amount of trypsin needed to activate aerolysin, we incubated pro-aerolysin with decreasing amounts of trypsin (Supplementary Fig S10B). The minimal dose of trypsin needed to activate aerolysin was chosen. Notably no oligomerization of aerolysin was observed during trypsin activation (Supplementary Fig S10B). We used HU/mL to normalize toxin activities in each experiment and to achieve consistent cytotoxicity across toxin preparations. To measure batch-to-batch variation, the LC_50_ were plotted by toxin type for toxin challenge with wild type cells (Supplementary Fig S10C-D). The highest toxin concentration that killed <20% of target cells by live cell imaging (e.g. TOPRO uptake) was defined as the sublytic dose (Ray et al., 2018). For live cell imaging experiments in HeLa cells, the sublytic dose used was 250 HU/mL for SLO, 250 HU/mL for ILY and 62 HU/mL for aerolysin. SLO ML was fluorescently labeled as previously described (Romero et al., 2017) while pro-aerolysin*^K244C^* was fluorescently labeled using maleimide chemistry. Briefly, purified Cys-less SLO ML was gel filtered into 100 mM sodium bicarbonate (pH 8.5) using a Zeba gel filtration column according to the manufacturer’s instructions. Sufficient monoreactive Cy5 dye (Cytiva) to label 0.25 mg protein was reconstituted in 100 mM sodium bicarbonate, added to the SLO ML, and incubated for 1 h at room temperature. Any unconjugated dye was removed by gel filtration of conjugated SLO ML into PBS. For the conjugation of Alexa Fluor 647 (AF647) to pro-aerolysin*^K244C^*, the toxin was eluted without DTT, then reduced with 6.32 mM Tris(2-carboxyethyl) phosphine hydrochloride for 30 min on ice. Alexa Fluor 647 maleimide (Sigma) dye was reconstituted as 1 µmol in DMSO and added to pro-aerolysin as 10-20 fold molar mass excess and incubated overnight at 4°C. Unconjugated dye was removed by gel filtration of conjugated pro-aerolysin*^K244C^* into Tris-salt buffer (50 mM Tris, 300 mM NaCl, pH 8). 5 mM DTT was added to the toxin, which was snap frozen on dry ice.

**Table 1.**
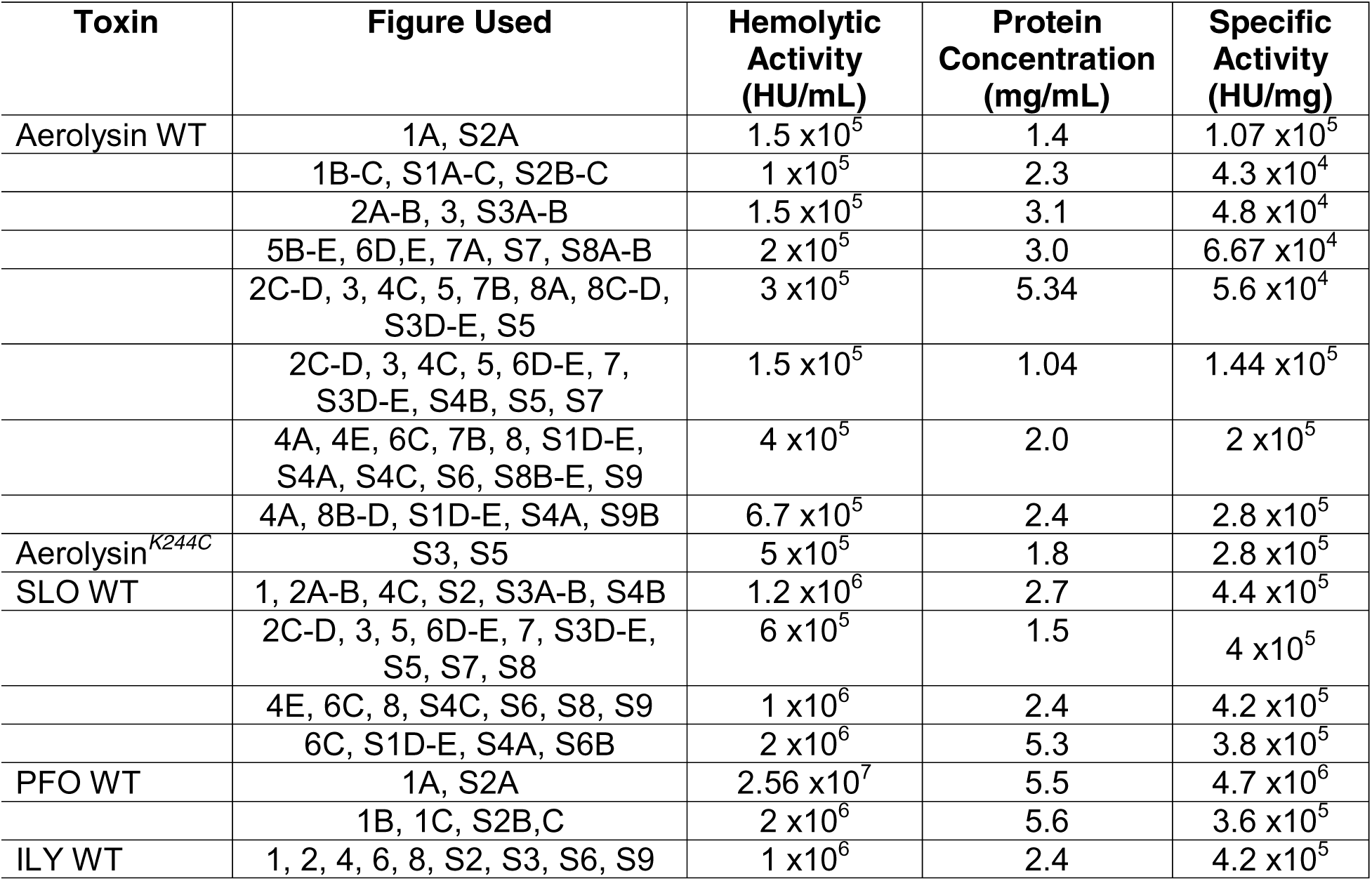
Specific activity of active toxin preps

### Trypsin activation of pro-aerolysin

Activation of pro-aerolysin was performed as previously described (Iacovache et al., 2011), with the following changes. Pro-aerolysin was activated by adding 1 µL of 0.02% trypsin in nuclease free water to 10 µL pro-aerolysin and incubating at RT for 10 minutes. The trypsin dilution was determined empirically based on the extent of pro-aerolysin cleavage.

### Transfection

HeLa cells were plated at 2 × 10^5^ cells per 35 mm glass-bottom dish or per well of a 6 well plate and transfected with 750 ng peGFP-N1 or GFP-Dysferlin using Lipofectamine

2000 in Opti-MEM two days before imaging or cytotoxicity assays. D10 media was replaced on the day after transfection. Transfection efficiencies for each construct ranged between 65-80%. For RNAi, HeLa cells were plated at 2 × 10^5^ cells in 6-well plates and transfected the following day with 10 nM siRNA using Lipofectamine 2000 in Opti-MEM for 48-72 h as previously described (Ray et al., 2022).

### MTT Assay

The MTT assay was performed and analyzed as previously described (Ray et al., 2018). Briefly, 2 x 10^4^ HeLa cells per well were plated in a 96 well tissue culture plate. On the next day, the media was aspirated. Cells were challenged with aerolysin (31-2000 HU/mL) for 1, 3, 6, 12, or 24 h at 37°C, then centrifuged at 1200× *g* for 5 min at 4°C to remove the media. Phenol Red free DMEM with 1.1 mM 3-(4,5-dimethylthiazol-2-yl)-2,5-diphenyltetrazolium bromide (MTT) was added, and cells were incubated at 37 °C for 4 h. SDS-HCl was used to dissolve formazan at 37°C overnight and absorbance was measured at 570 nm using a plate reader (Bio-Tek, Winooski, VT, USA). We calculated % viable cells as: % Viable = (Sample − Background)/ (Control − Background) × 100. Then, % Specific Lysis was calculated as 100 −%Viable.

### Flow Cytometry Assay

Cytotoxicity was performed as described (Haram et al., 2022; Keyel et al., 2012a; Ray et al., 2022; Ray et al., 2018; Romero et al., 2017). Briefly, 1×10^5^ cells were challenged in suspension with various concentrations of toxins for 5, 15, 30, or 60 min at 37°C in RPMI supplemented with 2 mM CaCl_2_ (RC) and 20 μg/mL PI. Cells were analyzed on a 4-laser Attune Nxt flow cytometer. Assay variations included changes to the media (replacing 2 mM CaCl_2_ with 2 mM EGTA or adding 150 mM KCl) or serum-starvation of cells in DMEM for 30 min at 37°C and pretreatment with DMSO or inhibitors in serum-free media for 30 min at 37 °C. For analysis of cell lysis, we gated out the debris and then quantified the percentage of cells with high dye fluorescence (2–3 log shift) (dye high), low dye fluorescence (∼1 log shift) (dye low), or background dye fluorescence (dye negative). Populations of cells marked “dye negative” and “dye low” remain metabolically active indicating that only the “dye high” population are dead cells) (Keyel et al., 2011). We calculated specific lysis as: % Specific Lysis = (% Dye High^Experimental^ − % Dye High^Control^)/ (100 − % Dye High^Control^) × 100. Transiently permeabilized cells were similarly calculated by using the “dye low” population instead of the “dye high” populations. The toxin dose needed to kill 50% of cells was defined as the Lethal Concentration 50% (LC_50_) and was determined by linear regression of the linear portion of the kill curve using Excel (Microsoft, Redmond, WA, USA) as previously described (Ray et al., 2018). The sublytic dose was defined as the highest toxin concentration that gave <20% specific lysis.

### Calcium flux assays

HeLa cells were plated at 2 × 10^5^ cells per 35 mm glass-bottom dish two days before imaging. Cells were labeled with 4 μM Fluo-4 AM (Cat# F14201) for 30 min at 37°C. After a brief wash, Fluo-4 AM labeled cells were challenged with the following sublytic toxins: 250 HU/mL SLO, 62.5 HU/ml Aerolysin or a mass equivalent of Aero*^Y221G^*, and imaged at 37°C in RPMI, 25 mM HEPES pH 7.4, 2 mM CaCl_2_, and 2 μg/mL TO-PRO3 for 45 min using a Fluoview 3000 confocal microscope (Olympus, Tokyo, Japan) equipped with a 60×, 1.42 NA oil immersion objective and recorded using the resonant scanner at 1-3 sec per frame. Fluo-4 AM was excited using the 488 nm laser, while TO-PRO3 was excited with the 640 nm laser. After 45 min, the assay was stopped by adding an equal volume of 2% Triton-X-100 to give a final concentration of 1%. Triton addition served as a control for TO-PRO uptake and dye loss. The extent of Ca^2+^ flux in cells was assessed by measuring intensity from the middle of the cell over time by plotting Z profiles in FIJI (ImageJ, NIH, Bethesda, MD, USA). TO-PRO3 uptake was determined in cells by plotting the Z profile of a nuclear subsample over time. We calculated fluorescence intensity over time for individual cells in each toxin group and then normalized each group to their brightest point to measure the temporal changes. We calculated the number of dead cells in each time point per experiment by converting monochromatic blue (TO-PRO-3) channel into masks and analyzing particles in FIJI. Experimental cells which expressed at least 2/3rd maximal fluorescence intensity of TO-PRO3 (post Triton-X-100) were considered dead. We categorized cells based on their survival: “died in 5 min”, “died between 5 and 45 min”, and “survived >45 min”.

### Ca^2+^ Oscillation Quantitation

To measure Ca^2+^ oscillations, we used realized volatility. Realized volatility measures the variance over a defined time period. We chose 10 sec periods using Nyquist’s theorem. We calculated the difference in the log fluorescence intensity at time t and time t+10, where t is in seconds. Then, we summed the square of these differences, and took the square root of this sum to get the final realized volatility.

### Fluorescent annexin shedding assays

HeLa cells were plated at 2 × 10^5^ cells per 35 mm glass-bottom dish and transfected with 750 ng A6-YFP using Lipofectamine2000 two days prior to imaging. The transfection efficiency was ∼65-80%. Transfected cells were challenged with sublytic toxin dose for SLO (250 HU/mL), aerolysin (62.5 HU/mL) or mass equivalent of aerolysin*^Y221G^* and imaged at 37°C in RPMI, 25 mM HEPES pH 7.4, and 2 mM CaCl_2_ with 2 μg/mL TO-PRO3 for 45 min using a Yokogawa CSU-X spinning disc confocal microscope (Intelligent Imaging Innovation, Denver, CO, USA). A6-YFP was excited using a 488 nm laser, while TO-PRO3 was excited with a 640 nm laser. Fluorescence was collected using a 60×, 1.49 NA oil immersion objective and recorded using an Evolve 512 EMCCD camera (Photometrics, Tucson, AZ, USA) at 1–3 sec/frame. After 45 min, an equal volume of 2% Triton-X-100 was added to give a final concentration of 1%. The total number of A6-YFP^+^ microvesicles released was counted manually using every second frame and was expressed as number of microvesicles shed/number of cells/minute. The percentage of cells showing A6-YFP translocation was determined by comparing the A6-YFP intensity at 15 min to the intensity 1 min after toxin addition. If these values were below 80% of the initial value, cells were considered to show A6-YFP translocation. In cells showing A6-YFP translocation, the extent of A6-YFP depletion from cells was assessed by measuring A6-YFP intensity from the middle of the cell over time by plotting Z profiles in ImageJ (NIH, Bethesda, MD, USA). TO-PRO3 uptake was determined in A6-YFP translocated cells by plotting the Z profile of a nuclear subsample over time. For both A6-YFP and TO-PRO3, the data were normalized to the integrated intensity of the brightest point. The t_1/2_ for annexin cytosolic depletion, membrane accumulation, or TO-PRO3 uptake was calculated by measuring the half-maximal intensity between the starting intensity and the average intensity of the last 4 min prior to Triton-X-100 addition. Membrane accumulation of A6 over time was calculated by determining the changes in the fluorescence intensity along the edges of the cells of translocated cells by plotting the Z profile in FIJI. The intensity was then normalized on a per-cell basis and individually analyzed cells were plotted. From the multi-plane images, several cells (>20) were analyzed from at least three independent experiments and graphed using Microsoft Excel and quantified values for individual cells were represented with Prism 8.1 (GraphPad, San Diego, CA, USA).

### Bleach correction and supplemental videos

Images were exported as .tif and split into individual monochromatic red, green, and blue channels in FIJI. For display (but not analysis), bleach correction was performed as previously described (Ray et al., 2018) on the “green” channel cells by histogram matching using FIJI followed by a median pass filter. The monochromatic images were then merged to form RGB .tif files, time-stamped, annotated, and exported as AVIs for supplementary videos.

### Immunofluorescence

Immunofluorescence was performed as previously described (Keyel et al., 2011). Briefly, HeLa cells (1 x 10^5^) were plated on coverslips, transfected with 750 ng of annexin A6-YFP and challenged with aerolysin or mass equivalent of aerolysin*^Y221G^* diluted in RPMI supplemented with 2 mM CaCl_2_ (RC) for 0, 15, 30, or 60 min at 37°C. Cells were washed in PBS, fixed in 2% paraformaldehyde, washed, permeabilized in 10% goat serum with 0.2% saponin, probed with rabbit polyclonal anti-aerolysin (1:500) for 1 h, washed, probed with goat Alexa-Fluor-647-conjugated anti-rabbit-IgG (1:1000) for 1 h, washed, DAPI stained, washed and mounted on slides in gelvatol. Cells were imaged on a Fluoview 3000 confocal microscope (Olympus, Tokyo, Japan) equipped with a 60×, 1.42 NA oil immersion objective. Images were processed using ImageJ (NIH, Bethesda, MD, USA).

### Isolation of Microvesicles (MVs)

MV were isolated as previously described (Keyel et al., 2011; Romero et al., 2017). Briefly, 5-10 × 10^6^ cells were harvested, resuspended in RC at 2.5-5 × 10^6^ cells/mL, challenged with a sublytic dose of toxin (hemolytic toxins), equivalent mass (inactive toxins), or 0.02% trypsin (control), and incubated for 15 min at 37°C. Challenged cells were then pelleted at 2000 × g for 10 min, solubilized at 95°C in SDS-sample buffer for 5 min, and sonicated. Supernatants were spun at 100,000 × g in a Beckman Coulter Optima XPN-80 ultracentrifuge using a Beckman SW 41 Ti rotor for 40 min at 4°C. The MV pellet was solubilized at 95°C in 4x SDS-PAGE sample buffer. We previously demonstrated (Keyel et al., 2011; Keyel et al., 2012b) that vesicles triggered by CDCs best fit the definition of MVs. These MV are nontoxic to cells and do not fuse with the plasma membrane of cells that internalize these MV (Keyel et al., 2012b).

### SDS-PAGE and immunoblotting

Samples were resolved on 10% polyacrylamide gels at 170 V/90 min and transferred to nitrocellulose in an ice bath with transfer buffer (15.6 mM Tris and 120 mM glycine) at 110 V for 90 min. Blots were blocked using 5% skim milk in 10 mM Tris-HCl, 150 mM NaCl and 0.1% Tween 20 at pH 7.5 (TBST), and portions of the blot were incubated with one of the following primary antibodies for 2 h in 1% skim milk in TBST at RT: 6D11 anti-SLO mAb (1:1000), CPTC-A1–3 anti-annexin A1 (1:250) mAb, anti-Annexin A2 (1:1000) mAb, anti-Alix (1:250) mAb, MANLAC-4A7 anti-Lamin A/C (1:250) mAb, AC-15 anti-β-actin (1:5000) or with rabbit polyclonal antibodies: anti-aerolysin, anti-annexin A6, anti-MEK, anti-phospho-MEK [Ser217/221], anti-ERK p44/42, anti-phospho-ERK [T202/Y404], anti-p38, or anti-phospho-p38, each at 1:1000 dilution. Next, blots were incubated with HRP-conjugated anti-mouse or anti-rabbit IgG antibodies (1:10,000) for 1 h in 1% skim milk in TBST at RT and developed with ECL: either 0.01% H_2_O_2_ (Walmart, Fayetteville, AR, USA), 0.2 mM p-Coumaric acid (Sigma), mM Luminol (Sigma) in 0.1 M Tris pH 8.4, ECL Plus Western Blotting Substrate (Cat# 32134), ECL Prime Western Blotting Reagent (Cat# RPN2232, Cytiva), or Pierce ECL western blotting substrate (Lot # UH288963A, Thermo Fisher).

### ELISA

ELISA plates were coated with 10 ng of pro-aerolysin, pro-aerolysin*^Y221G^*, aerolysin or aerolysin*^Y221G^* and incubated overnight at 4°C. Plates were then washed 3 times in PBS with 0.05% Tween-20 (PBST), blocked for 1 h with 1% BSA in PBST, and then serially diluted pre-immune serum, the affinity-purified anti-aerolysin antibody, or the depleted serum post-affinity purification were added for 2 h at 37°C. Plates were washed 3x in PBST, incubated with HRP-conjugated goat anti-rabbit IgG (1:20,000) and developed using 0.2 mg/mL TMB (Sigma), 0.015% H_2_O_2_ in 100 mM sodium acetate, pH 5.5. The reaction was stopped with 0.5 M H_2_SO_4_. A_450_ was measured on a plate reader (Bio-tek, Winooski, VT) and plotted against antibody dilution.

### Statistics

Prism 8.1 (GraphPad, San Diego, CA, USA) or Excel were used for statistical analysis. Data are represented as mean ± SEM as indicated. The LC_50_ for toxins was calculated by logistic modeling in Excel as previously described (Haram et al., 2022). Statistical significance was determined by regular or repeated measures one-way or two-way ANOVA as specified; p < 0.05 was considered to be statistically significant. Graphs were generated in Excel, Prism and Photoshop (Adobe, San Jose, CA, USA).

## Acknowledgments

The authors would like to thank members of the Keyel lab for critical review of the manuscript. We thank colleagues for the generous gifts of reagents. We thank the College of Arts & Sciences Microscopy for use of facilities. This work was supported by American Heart Association grant 16SDG30200001, Texas Tech University, and NIH grant 1R21AI156225 to PAK.

## Author Contributions

Conceptualization: PAK

Methodology: RT, PAK

Investigation: RT, PAK

Data Curation: RT, PAK

Formal Analysis: RT, PAK

Supervision: PAK

Project Administration: PAK

Funding Acquisition: PAK

Writing—original draft: RT, PAK

Writing—review & editing: RT, PAK

## Data and materials availability

All data are available in the main text or the supplementary materials.

## Figure Legends

## Supplemental Figure Legends

**Supplementary Figure S1. Dose and time affect the aerolysin activity.** (A-C) HeLa cells were unchallenged or challenged with indicated concentrations of aerolysin (Aero), mutant aerolysin^Y221G^ (Aero*^Y221G^*) or trypsin in 2 mM CaCl_2_ supplemented RPMI (RC) with 20 μg/ml PI for 5-60 min at 37°C. PI uptake was analyzed by flow cytometry. (A) The gating strategy for flow cytometric cytotoxicity assay is shown. (B, C) Time and dose dependent cytotoxic responses are shown. (D, E) HeLa cells were challenged with 31-2000 HU/mL aerolysin in RC for 1-24 h. Viability was measured by MTT assay. Specific lysis was determined and LC_50_ was calculated as described in methods. Graphs display the (B, C, E) mean ± SEM of 3 independent experiments or (D) median of 5 individual experiments. For (D) data points represent individual experiments.

**Supplementary Figure S2. Aerolysin is more toxic to cells at lower activities than CDCs.** (A) HeLa, (B) 3T3, or (C) Caspase 1/11^-/-^ bone-marrow derived macrophages (MΦ) were challenged with the indicated concentrations of (A) ILY, or (A-C) pro-aerolysin (Pro-Aero), aerolysin (Aero), SLO, PFO, or a mass equivalent to wild type aerolysin of aerolysin^Y221G^ (Aero*^Y221G^*), in RC medium with 20 μg/ml PI for 30 min at 37°C. cflow cytometry. Specific lysis was determined. Graphs display the mean ± SEM of (A) 8 or (B, C) 6 independent experiments.

**Supplementary Figure S3. Removal of Ca^2+^ influx protects cells against aerolysin.** (A) HeLa cells were challenged with 31-2000 HU/mL pro-aerolysin (Pro-Aero), aerolysin (Aero), or SLO with or without 150 mM KCl. PI uptake was analyzed by flow cytometry. (B) HeLa cells were challenged with 31-2000 HU/mL pro-aerolysin, aerolysin, SLO, or a mass equivalent to aerolysin of mutant aerolysin^Y221G^ (Aero*^Y221G^*) for 30 min at 37°C either in 2 mM CaCl_2_ or 2 mM EGTA supplemented RPMI. PI uptake was analyzed by flow cytometry. (C) HeLa cells were challenged with AlexaFluor647-labeled pro-aerolysin^K244C^ (Pro-aero*^K244C^*) at the indicated concentrations either in 2 mM CaCl_2_ or 2 mM EGTA supplemented RPMI. Toxin binding was analyzed by flow cytometry. Median fluorescent intensity (MFI) was calculated. (D) HeLa cells were challenged with 31-2000 HU/mL aerolysin, SLO, or ILY for 30 min at 37°C in 2 mM CaCl_2_ supplemented RPMI or Tyrode’s buffer. PI uptake was analyzed by flow cytometry. (E) The individual time to death for the aerolysin- and SLO-challenged cells in Fig 3 were plotted. (F) Realized Volatility of Ca^2+^ flux was calculated as described in methods. Graphs show the average ± SEM of at least (A, B) 5, (C) 3, or (D) 6 independent experiments. (F) Violin plot summarizes individual cell data from 3 independent experiments. Letters (a-c) denote statistically significant (p < 0.05) groups using repeated-measures ANOVA between groups.

**Supplementary Figure S4. MAP kinase signaling does not protect cells from proximal aerolysin damage.** (A-C) HeLa cells were serum starved for 30 min and pre-treated with DMSO, 20 μM SB203580 (p38i), or 20 μM SP600125 (JNKi) for 30 min and challenged either in 2 mM CaCl_2_ or 2 mM EGTA supplemented RPMI with 31-2000 HU/mL aerolysin, SLO, or ILY for 30 min at 37°C. PI uptake was analyzed by flow cytometry. Specific lysis was determined. Graphs show the average ± SEM of at least (A) 6, (B) 7, or (C) 5 independent experiments.

**Supplementary Figure S5. Cavin-1 deficiency and adipocyte differentiation do not prevent toxin binding to cells.** (A) Undifferentiated wild type (WT), K5, or K3 3T3-L1 cells were challenged with 31-2000 HU/mL aerolysin or SLO in RC medium for 30 min at 37°C. PI uptake was analyzed by flow cytometry. Specific lysis was determined. (B-E) WT, K5, or K3 3T3-L1 cells were differentiated to adipocytes and collected at (B-C) 9 or (D-E) 14 days. HeLa cells and adipocytes were challenged with (B, D) AlexaFluor 647-labeled pro-aerolysin*^K244C^* (Pro-Aero*^K244C^*) or (C, E) Cy5-labeled monomer-locked SLO (SLO ML) at the indicated concentrations in RC with 20 μg/mL Pl for 30 min at 4°C. Toxin binding was analyzed by flow cytometry after gating cells by viability using PI staining. Graphs show the average ± SEM of (A) 7 or (B-E) 4 independent experiments.

**Supplementary Figure S6. Annexins minimally resist aerolysin.** HeLa cells were transfected with siRNAs to control (CTL) or annexins for 72 h. Cells were then challenged with 31-2000 HU/mL (A) aerolysin (Aero), (B) SLO, or (C) ILY for 30 min at 37°C. PI uptake was analyzed by flow cytometry. Specific lysis was determined as described in the methods. Graphs show the average ± SEM of 4 independent experiments.

**Supplementary Figure S7. Aerolysin provokes slower annexin A6 translocation compared to SLO.** A6-YFP transfected HeLa cells were challenged with sublytic concentrations of toxins (62 HU/mL pro-aerolysin (Pro-Aero) or aerolysin (Aero), or 250 HU/mL SLO), or mass equivalent to aerolysin of mutant aerolysin^Y221G^ (Aero*^Y221G^*) in imaging buffer. Cells were imaged by confocal microscopy for ∼45 min at 37°C and then lysed with 1% Triton. The average (A) annexin depletion from the cytosol, (B) annexin enrichment on the membrane, or (C) TO-PRO3 uptake over time was normalized to maximal intensity and graphed. For each toxin, 25–30 cells from 3 independent experiments were analyzed. (D) A replicate blot from Fig 7 was probed with anti-aerolysin. * indicates non-specific bands.

**Supplementary Figure S8. The affinity-purified polyclonal aerolysin antibody is specific to aerolysin.** (A) 20 µg of pro-aerolysin (Pro-Aero), pro-aerolysin^Y221G^ (Aero*^Y221G^*), aerolysin (Aero), or aerolysin^Y221G^ (Aero*^Y221G^*) were resolved by SDS-PAGE on a 10% polyacrylamide gel and transferred to nitrocellulose. The nitrocellulose was probed with the indicated dilutions of pre-immune and production bleed sera from two immunized rabbits. (B) Western blotting of 100 µg or 20 µg of pro-aerolysin and aerolysin was performed using the indicated dilutions the anti-aerolysin antibody after affinity purification. (C) HeLa cells were unchallenged or challenged with 125 or 1000 HU/mL aerolysin or SLO or a mass of mutant Aero*^Y221G^* equivalent to 1000 HU/mL aerolysin for 15 min at 37°C. Cells were lysed in 95°C 1x SDS buffer and sonicated. Western blots on cell lysates were performed using pre-immune serum, affinity purified anti-aerolysin antibody, 6D11 anti-SLO, or AC-15 anti-β-Actin antibody. (D) ELISA plates were coated with 10 ng of pro-aerolysin, pro-aerolysin*^Y221G^*, aerolysin or aerolysin*^Y221G^*. ELISA was performed using pre-immune serum, the affinity-purified anti-aerolysin antibody, or the depleted serum post-affinity purification, followed by HRP-conjugated anti-rabbit IgG and TMB. (E) A6-transfected (green) HeLa cells were either unchallenged (control) or challenged with 62, 125 and 1000 HU/mL aerolysin or mass equivalent to 1000 HU/mL aerolysin of mutant Aero*^Y221G^* for 15 min, then fixed, permeabilized and blocked. Then cells were probed with the affinity purified anti-aerolysin antibody (1:500 dilution) followed by goat anti-rabbit IgG conjugated to Alexa Fluor 647 (red). Nuclei were stained with DAPI (blue). Scale bar = 10 μm.

**Supplementary Figure S9. Dysferlin-mediated patch repair protects cells from aerolysin**. HeLa cells were either untransfected or transfected with GFP, or GFP-Dysferlin (GFP-Dysf) for 48 h and challenged with 31-2000 HU/mL aerolysin (Aero), SLO or ILY for 30 min at 37°C either in 2 mM CaCl_2_ or 2 mM EGTA supplemented RPMI. PI uptake was analyzed by flow cytometry. Graphs show the mean ± S.E.M of (A) 7, or (B) 5 independent experiments.

**Supplementary Figure S10. Purification and quality control of aerolysin minimizes batch to batch variation.** (A) Pro-aerolysin or pro-aerolysin*^Y221G^* were purified by size exclusion chromatography for antibody generation, and toxin assays. (B) His-purified pro-aerolysin was incubated with the indicated amount of trypsin for 10 min at room temp and analyzed by SDS-PAGE. The trypsin dose chosen was the lowest dose converting pro-aerolysin to aerolysin. Representative Coomassie-stained gels are shown. (C-D). The LC_50_ for wild type HeLa cells challenged with (C) aerolysin or (D) SLO in this manuscript are plotted against the specific activity of the toxin batch. High purity aerolysin preps are denoted by triangles (5.6 x 10^4^ HU/mg) or diamonds (14.4 x 10^4^ HU/mg). Aerolysin toxin prep was statistically significant by Kruskal-Wallis test. However Dunn multiple comparison post hoc testing only showed statistical significance between 14.4 x 10^4^ and 28 x 10^4^ HU/mg preps. There was no difference by toxin purity.

**Supplementary Figure S11. Full western blots for MEK activation**. Full western blots from Fig 4 are displayed.

**Supplementary Figure S12. Full western blots for microvesicle shedding**. Full western blots from Fig 7 are displayed.

## Supplemental Video Legends

**Supplemental Video V1. Aerolysin triggers Ca^2+^ flux.** HeLa cells were labeled with Fluo-4-AM (green) for 30 min in DMEM. The media was replaced with imaging buffer (2 μg/mL TO-PRO3 (blue), 25 mM HEPES, pH 7.4, and 2 mM CaCl_2_ supplemented RPMI). Cells were imaged by confocal microscopy at 1-3 frames/sec for ∼45 min at 37°C. Sublytic (A) SLO (250 HU/mL), or (B) aerolysin (62 HU/mL) was added 5 min after imaging started. Triton-X-100 was added just prior to the end of imaging as a positive control for cell permeabilization. Images were bleach-corrected by histogram matching. Time displays min post toxin challenge. Representative Ca^2+^-flux is shown by arrowheads. Scale bar = 10 μm.

**Supplemental Video V2. Ca^2+^ flux following control aerolysin challenge.** HeLa cells were labeled with Fluo-4-AM (green) for 30 min in DMEM. The media was replaced with imaging buffer supplemented either with 2 mM Ca^2+^ or 2 mM EGTA, including TO-PRO-3 (blue). Cells were imaged by confocal microscopy at 1-3 frames/sec for ∼45 min at 37°C. Sublytic (A) aerolysin (62 HU/mL) in presence of 2 mM EGTA or (B) mass equivalent to aerolysin of mutant Aero*^Y221G^*, or (C) PBS were added 5 min after imaging started. Triton-X-100 was added just prior to the end of imaging as a positive control for cell permeabilization. Images were bleach-corrected by histogram matching. Time displays min post toxin challenge. Scale bar = 10 μm.

**Supplemental Video V3. Intrinsic repair drives shedding of A6^+^ microvesicle following aerolysin challenge.** HeLa cells transfected with A6-YFP (green) were challenged with sublytic (A) SLO (250 HU/mL), (B) pro-aerolysin (62 HU/mL), (C) aerolysin (aero) (62 HU/mL), or (D) mass equivalent of aerolysin^Y221G^ (Aero*^Y221G^*) to aerolysin in imaging buffer including 2 μg/mL TO-PRO3 (blue) and imaged at 37°C by live cell confocal imaging at 1-3.5 frame/second. Triton-X-100 was added just prior to the end of imaging as a positive control for cell permeabilization. Images were bleach-corrected by histogram matching. Time displays min after toxin challenge. A6-YFP recruitment to the sites of plasma membrane damage are shown by arrowheads. Scale bar = 10 μm.

**Supplemental Video V4. GFP is not recruited to the membrane during toxin challenge.** GFP (green) transfected HeLa cells were challenged with sublytic (A) SLO (250 HU/mL), (B) ILY (250 HU/mL), (C) aerolysin (Aero) (62 HU/mL), or (D) mass equivalent to aerolysin of aerolysin^Y221G^ (AeroY221G) in imaging buffer including 2 μg/mL TO-PRO3 (blue) and imaged at 37°C by live cell confocal imaging at 1-3.5 frame/second. Triton-X-100 was added just prior to the end of imaging as a positive control for cell permeabilization. Images were bleach-corrected by histogram matching. Time displays min post toxin challenge. Scale bar = 5 μm.

**Supplemental Video V5. GFP-Dysferlin fuses with the plasma membrane during toxin challenge.** GFP-Dysf (green) transfected HeLa cells were challenged with sublytic (A) SLO (250 HU/mL), (B) ILY (250 HU/mL), (C) aerolysin (62 HU/mL), or (D) mass equivalent to aerolysin of aerolysin^Y221G^ (aero*^Y221G^*) in imaging buffer including 2 μg/mL TO-PRO3 (blue) and imaged at 37°C by live cell confocal imaging at 1-3.5 frame/second. Triton-X-100 was added just prior to the end of imaging as a positive control for cell permeabilization. Images were bleach-corrected by histogram matching. Time displays min post toxin challenge. Depletion of GFP-Dysf containing vesicles from cytosol are shown by arrowheads. Scale bar = 5 μm.

**Supplemental Video V6. Removal of extracellular calcium reduces patch repair.** GFP-Dysf (green) transfected HeLa cells were challenged with sublytic (A) SLO (250 HU/mL), (B) ILY (250 HU/mL), or (C) aerolysin (62 HU/mL) in imaging buffer including 2 μg/mL TO-PRO3 (blue) and imaged at 37°C by live cell confocal imaging at 1-3.5 frame/second. 2 mM EGTA was used instead of 2 mM CaCl_2_ in imaging buffer. Triton-X-100 was added just prior to the end of imaging as a positive control for cell permeabilization. Images were bleach-corrected by histogram matching. Time displays min post toxin challenge. Scale bar = 5 μm.

