## Supplemental Figures for "Patch repair protects cells from the small pore-forming toxin aerolysin"

### Figure S1

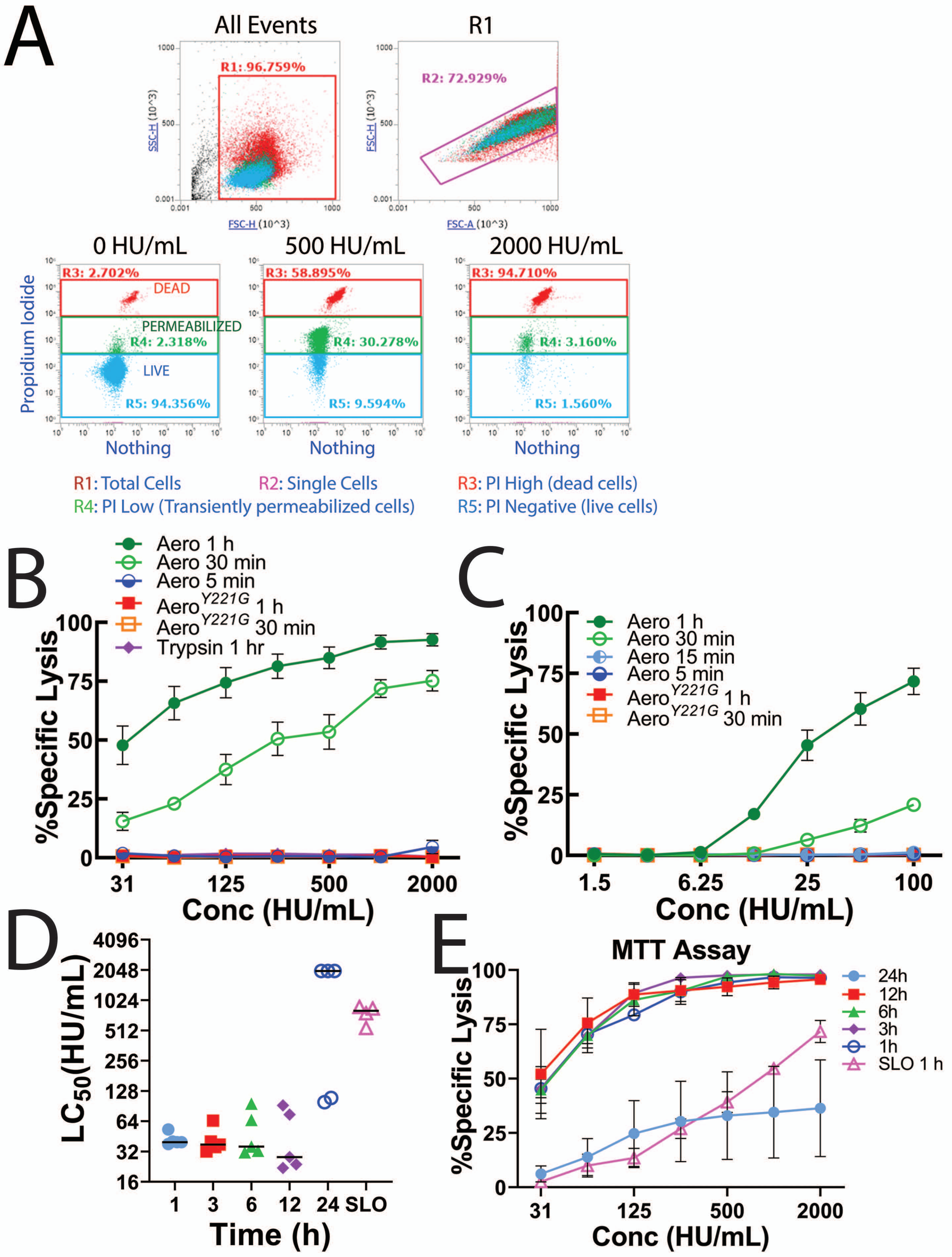

A
All Events
R1
R1: 96.759%
R2: 72.929%
SSC-H (10^3)
FSC-H (10^3)
FSC-H (10^3)
FSC-A (10^3)
0 HU/mL
500 HU/mL
2000 HU/mL
R3: 2.702%
R3: 58.895%
R3: 94.710%
DEAD
PERMEABILIZED
R4: 2.318%
R4: 30.278%
R4: 3.160%
LIVE
R5: 94.356%
R5: 9.594%
R5: 1.560%
Propidium Iodide
Nothing
R1: Total Cells
R2: Single Cells
R3: PI High (dead cells)
R4: PI Low (Transiently permeabilized cells)
R5: PI Negative (live cells)
B
Aero 1 h
Aero 30 min
Aero 5 min
Aero Y221G 1 h
Aero Y221G 30 min
Trypsin 1 hr
%Specific Lysis
Conc (HU/mL)
C
Aero 1 h
Aero 30 min
Aero 15 min
Aero 5 min
Aero Y221G 1 h
Aero Y221G 30 min
%Specific Lysis
Conc (HU/mL)
D
LC50(HU/mL)
Time (h)
1
3
6
12
24
SLO
E
MTT Assay
24h
12h
6h
3h
1h
SLO 1 h
%Specific Lysis
Conc (HU/mL)

### Figure S2

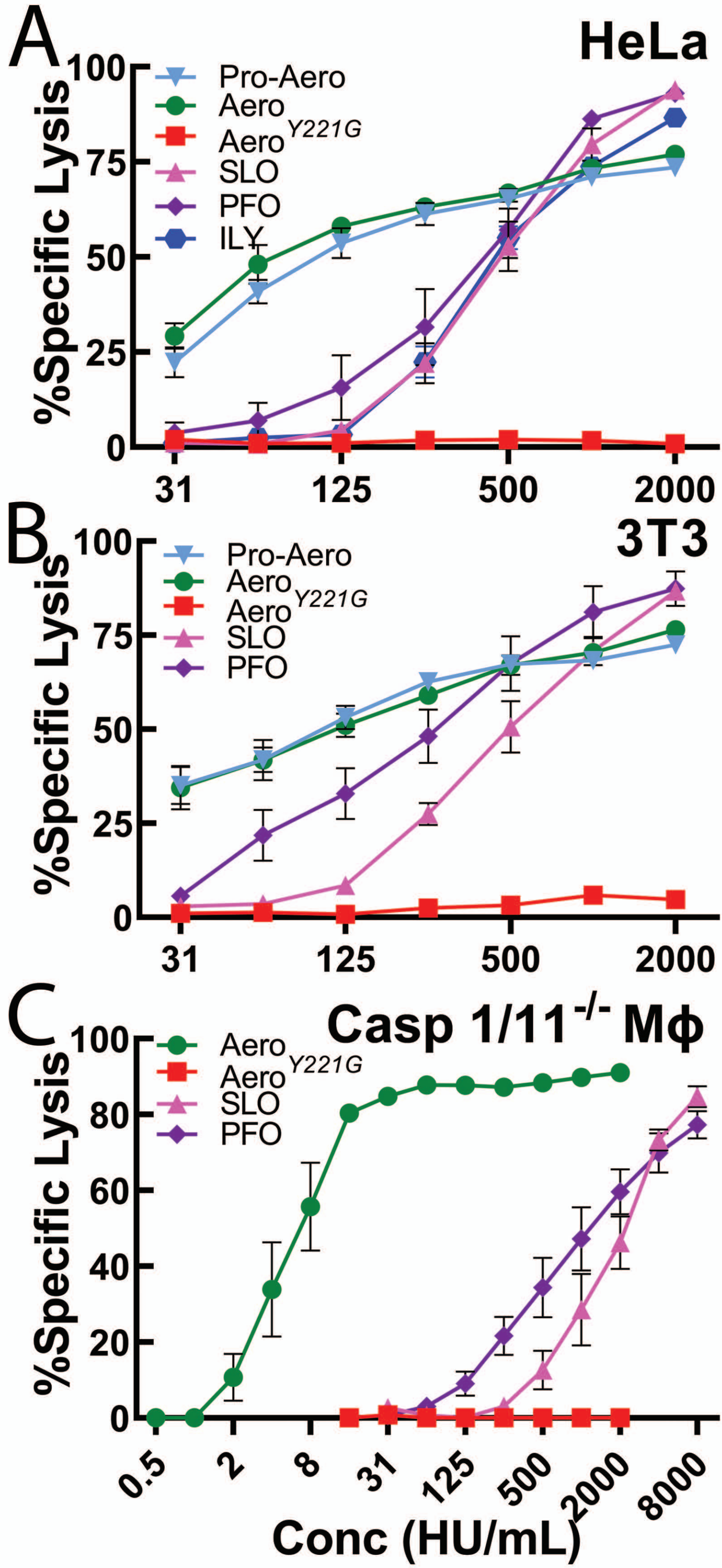
A
HeLa
%Specific Lysis
100
75
50
25
0
Pro-Aero
Aero
Aero Y221G
SLO
PFO
ILY
31
125
500
2000
B
3T3
%Specific Lysis
100
75
50
25
0
Pro-Aero
Aero
Aero Y221G
SLO
PFO
31
125
500
2000
C
Casp 1/11 -/- Mϕ
%Specific Lysis
100
80
60
40
20
0
Aero
Aero Y221G
SLO
PFO
0.5
2
8
31
125
500
2000
8000
Conc (HU/mL)

### Figure S3

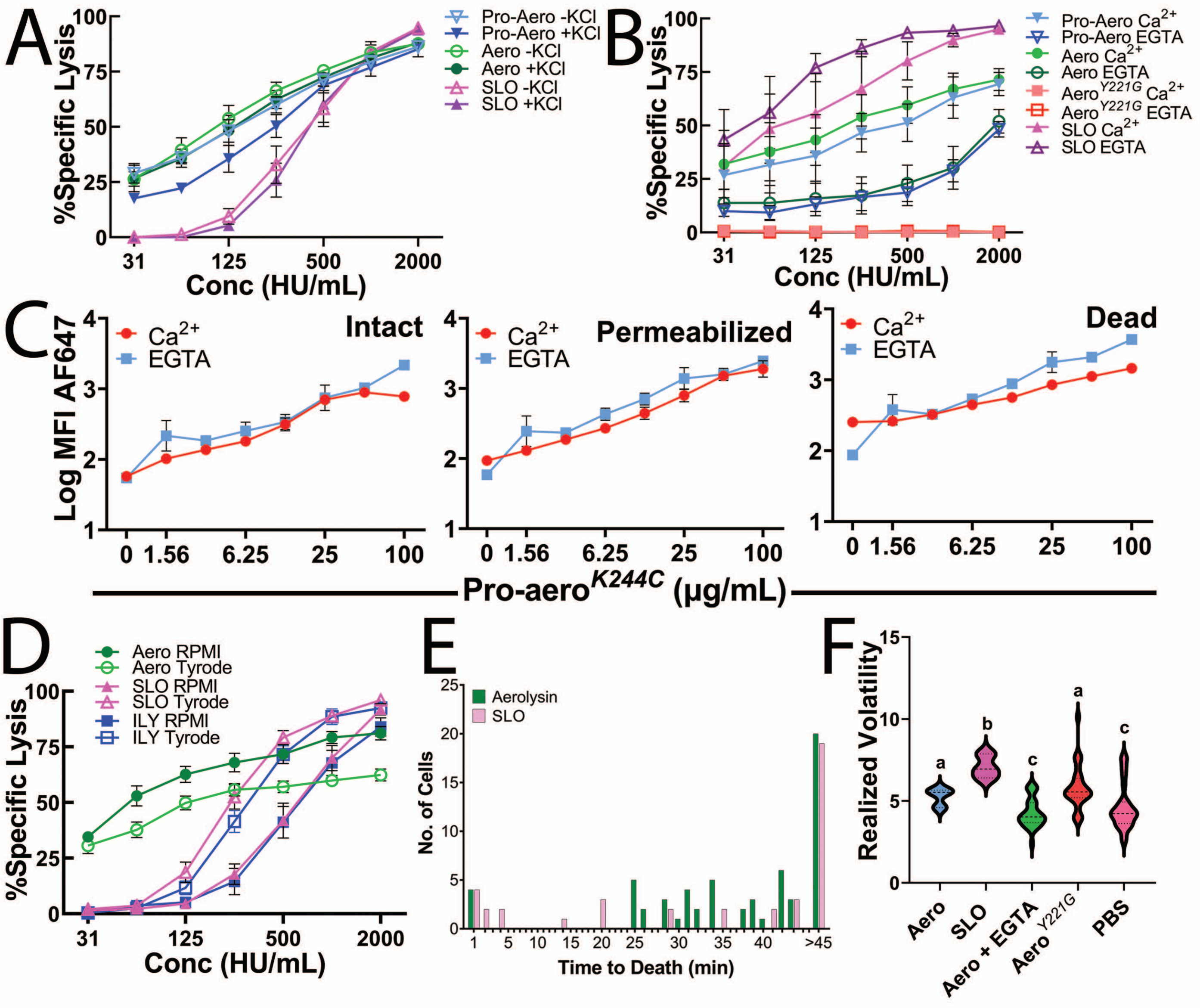
A
%Specific Lysis
Conc (HU/mL)
Pro-Aero -KCl
Pro-Aero +KCl
Aero -KCl
Aero +KCl
SLO -KCl
SLO +KCl
B
%Specific Lysis
Conc (HU/mL)
Pro-Aero Ca2+
Pro-Aero EGTA
Aero Ca2+
Aero EGTA
Aero Y221G Ca2+
Aero Y221G EGTA
SLO Ca2+
SLO EGTA
C
Log MFI AF647
Ca2+
EGTA
Intact
Permeabilized
Dead
Pro-aero K244C (µg/mL)
D
Aero RPMI
Aero Tyrode
SLO RPMI
SLO Tyrode
ILY RPMI
ILY Tyrode
%Specific Lysis
Conc (HU/mL)
E
Aerolysin
SLO
No. of Cells
Time to Death (min)
F
Realized Volatility
Aero
SLO
Aero + EGTA
Aero Y221G
PBS

### Figure S4

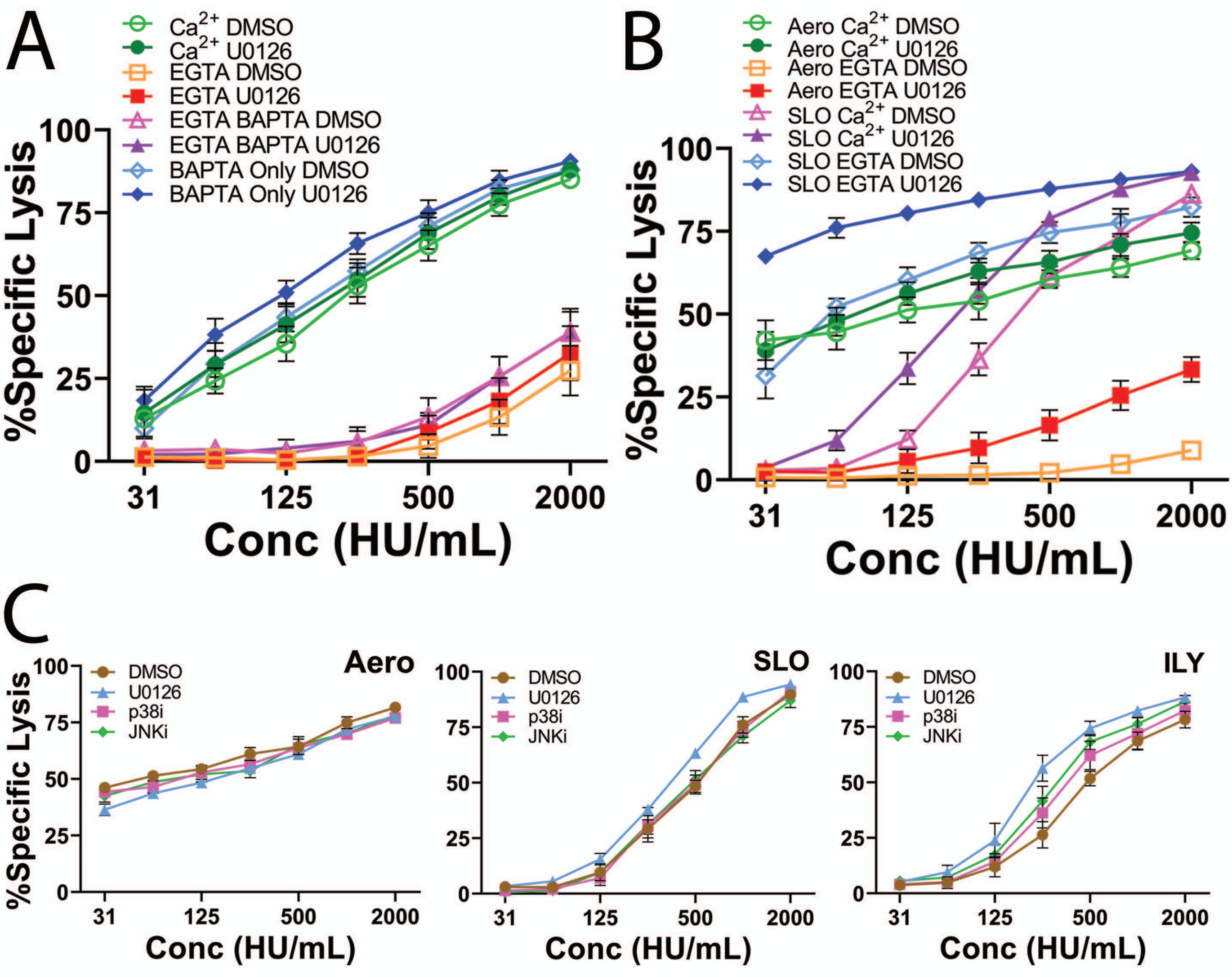
A
Ca2+ DMSO
Ca2+ U0126
EGTA DMSO
EGTA U0126
EGTA BAPTA DMSO
EGTA BAPTA U0126
BAPTA Only DMSO
BAPTA Only U0126
%Specific Lysis
Conc (HU/mL)
B
Aero Ca2+ DMSO
Aero Ca2+ U0126
Aero EGTA DMSO
Aero EGTA U0126
SLO Ca2+ DMSO
SLO Ca2+ U0126
SLO EGTA DMSO
SLO EGTA U0126
%Specific Lysis
Conc (HU/mL)
C
Aero
SLO
ILY
DMSO
U0126
p38i
JNKi
%Specific Lysis
Conc (HU/mL)
0
25
50
75
100
31
125
500
2000

### Figure S5

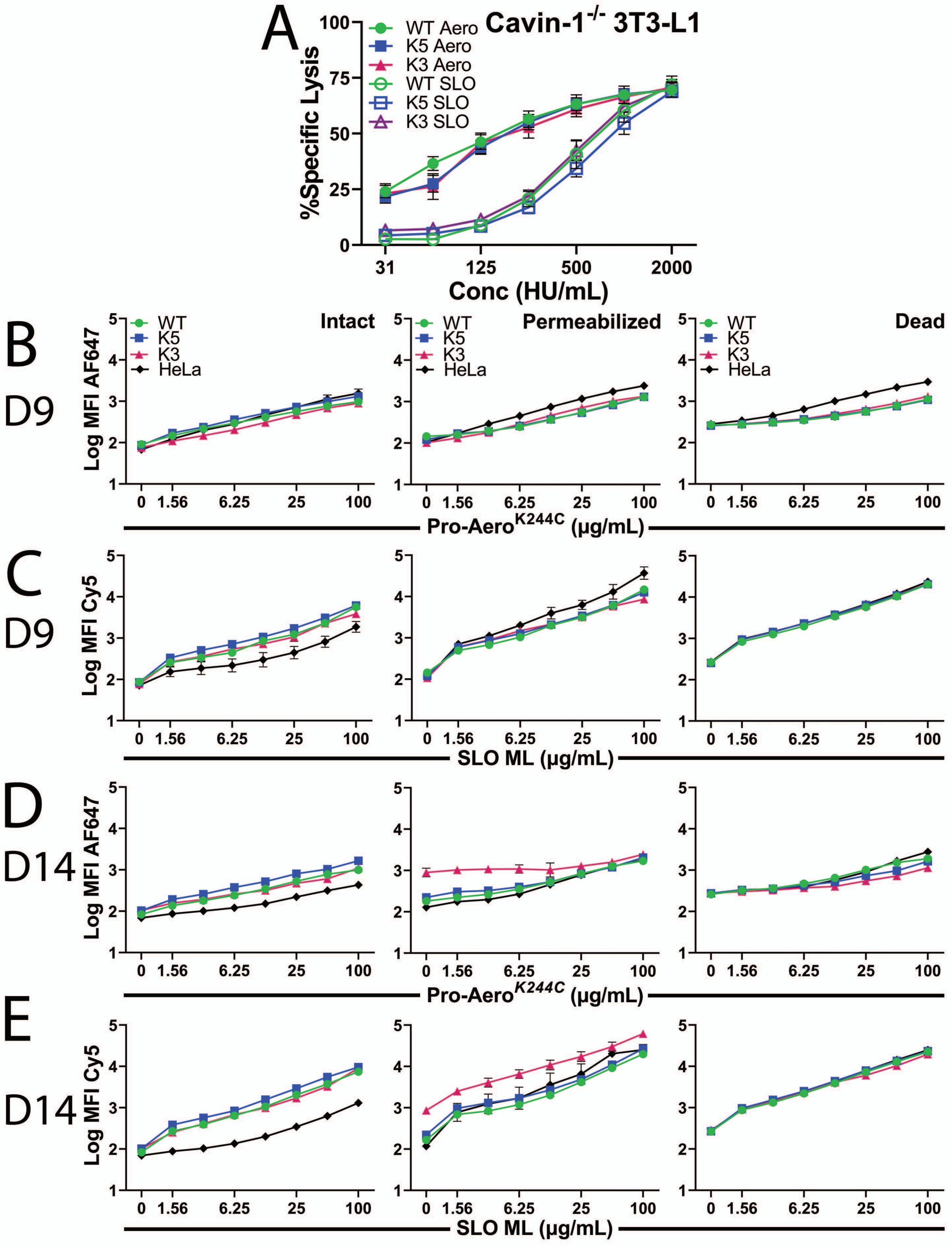
A
Cavin-1-/- 3T3-L1
%Specific Lysis
Conc (HU/mL)
WT Aero
K5 Aero
K3 Aero
WT SLO
K5 SLO
K3 SLO
B
D9
C
D9
D
D14
E
D14
Intact
Permeabilized
Dead
WT
K5
K3
HeLa
Log MFI AF647
Log MFI Cy5
Pro-Aero K244C (µg/mL)
SLO ML (µg/mL)

### Figure S6

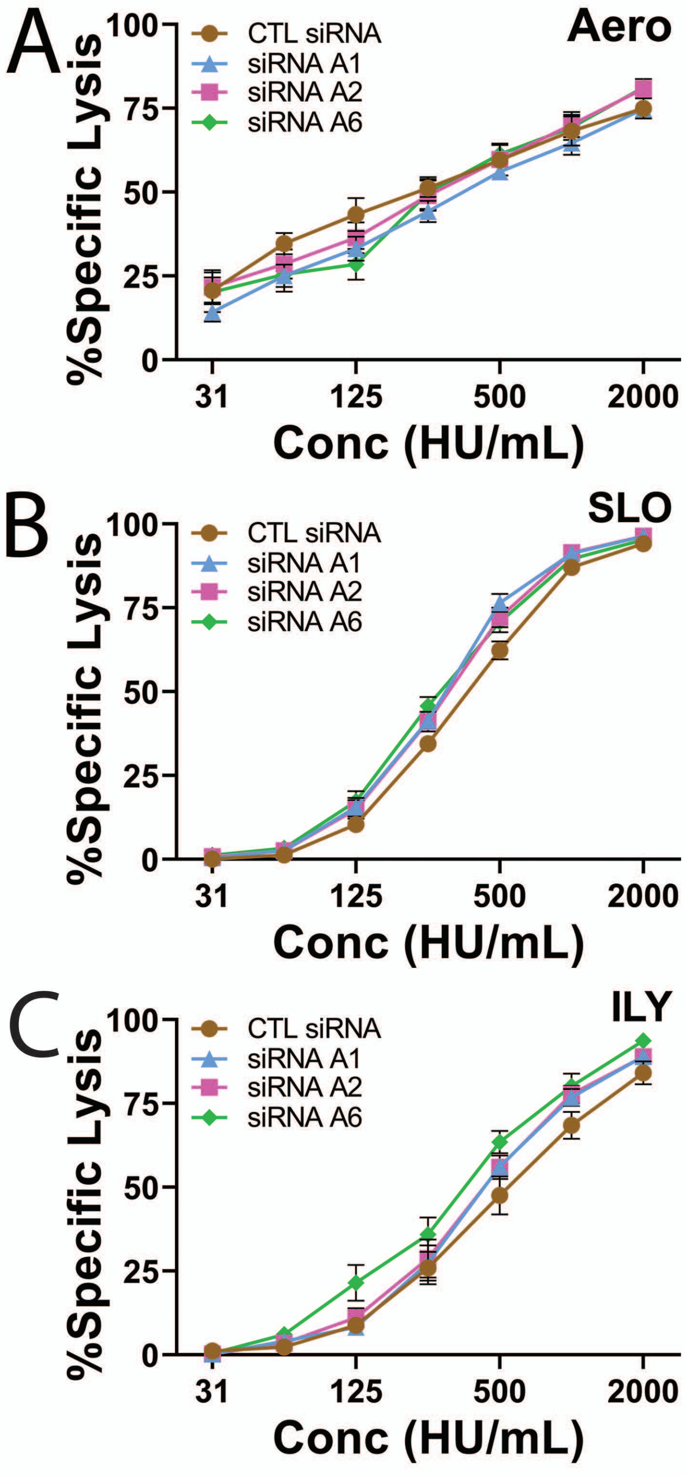
A
Aero
CTL siRNA
siRNA A1
siRNA A2
siRNA A6
%Specific Lysis
0
25
50
75
100
31
125
500
2000
Conc (HU/mL)
B
SLO
CTL siRNA
siRNA A1
siRNA A2
siRNA A6
%Specific Lysis
0
25
50
75
100
31
125
500
2000
Conc (HU/mL)
C
ILY
CTL siRNA
siRNA A1
siRNA A2
siRNA A6
%Specific Lysis
0
25
50
75
100
31
125
500
2000
Conc (HU/mL)

Figure S7

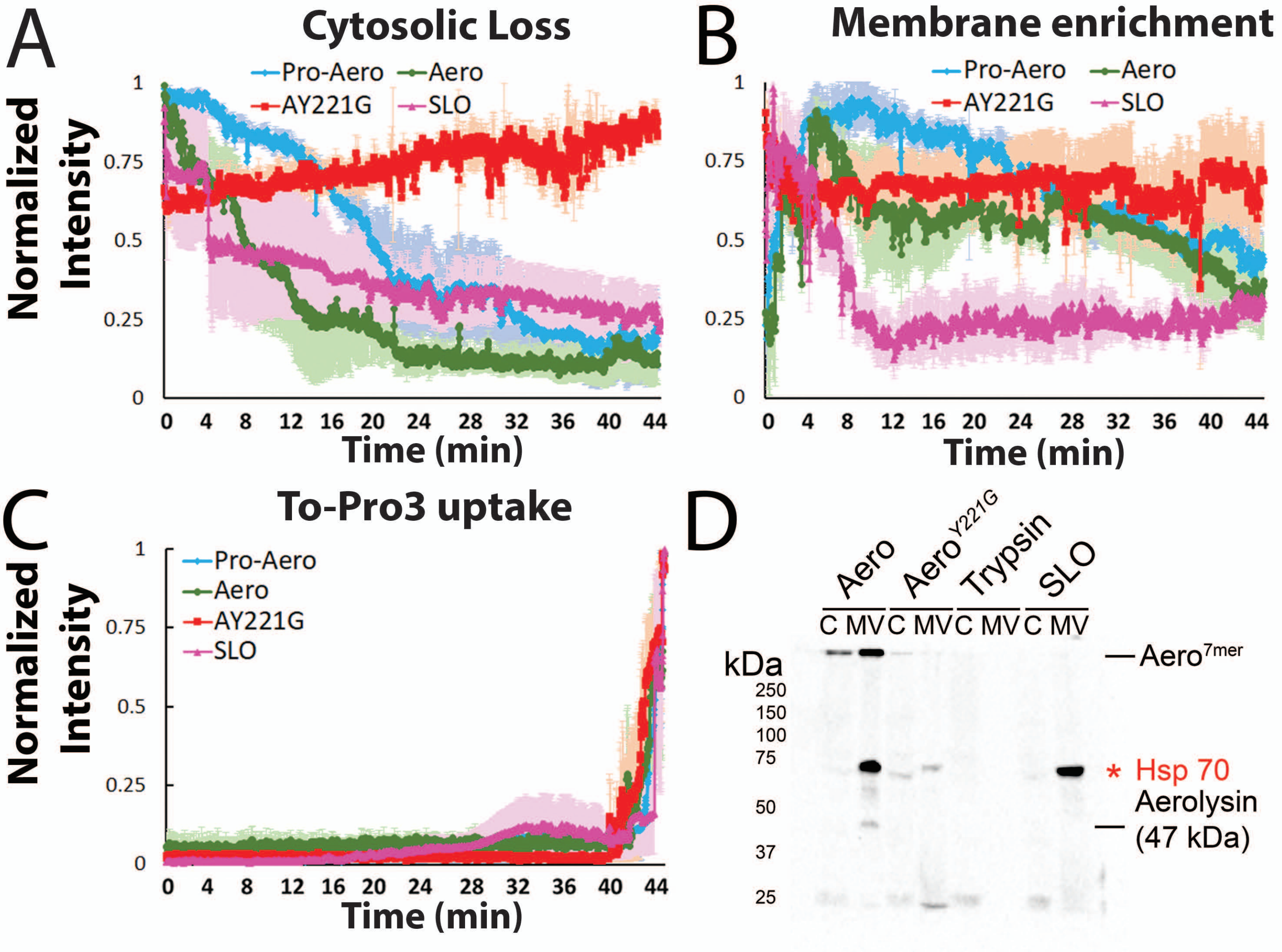

A
Cytosolic Loss
Pro-Aero
Aero
AY221G
SLO
Normalized Intensity
1
0.75
0.5
0.25
0
0 4 8 12 16 20 24 28 32 36 40 44
Time (min)
B
Membrane enrichment
Pro-Aero
Aero
AY221G
SLO
1
0.75
0.5
0.25
0
0 4 8 12 16 20 24 28 32 36 40 44
Time (min)
C
To-Pro3 uptake
Pro-Aero
Aero
AY221G
SLO
Normalized Intensity
1
0.75
0.5
0.25
0
0 4 8 12 16 20 24 28 32 36 40 44
Time (min)
D
Aero
Aero Y221G
Trypsin
SLO
C MV C MV C MV C MV
kDa
250
150
100
75
50
37
25
Aero 7mer
* Hsp 70
Aerolysin (47 kDa)

### Figure S8

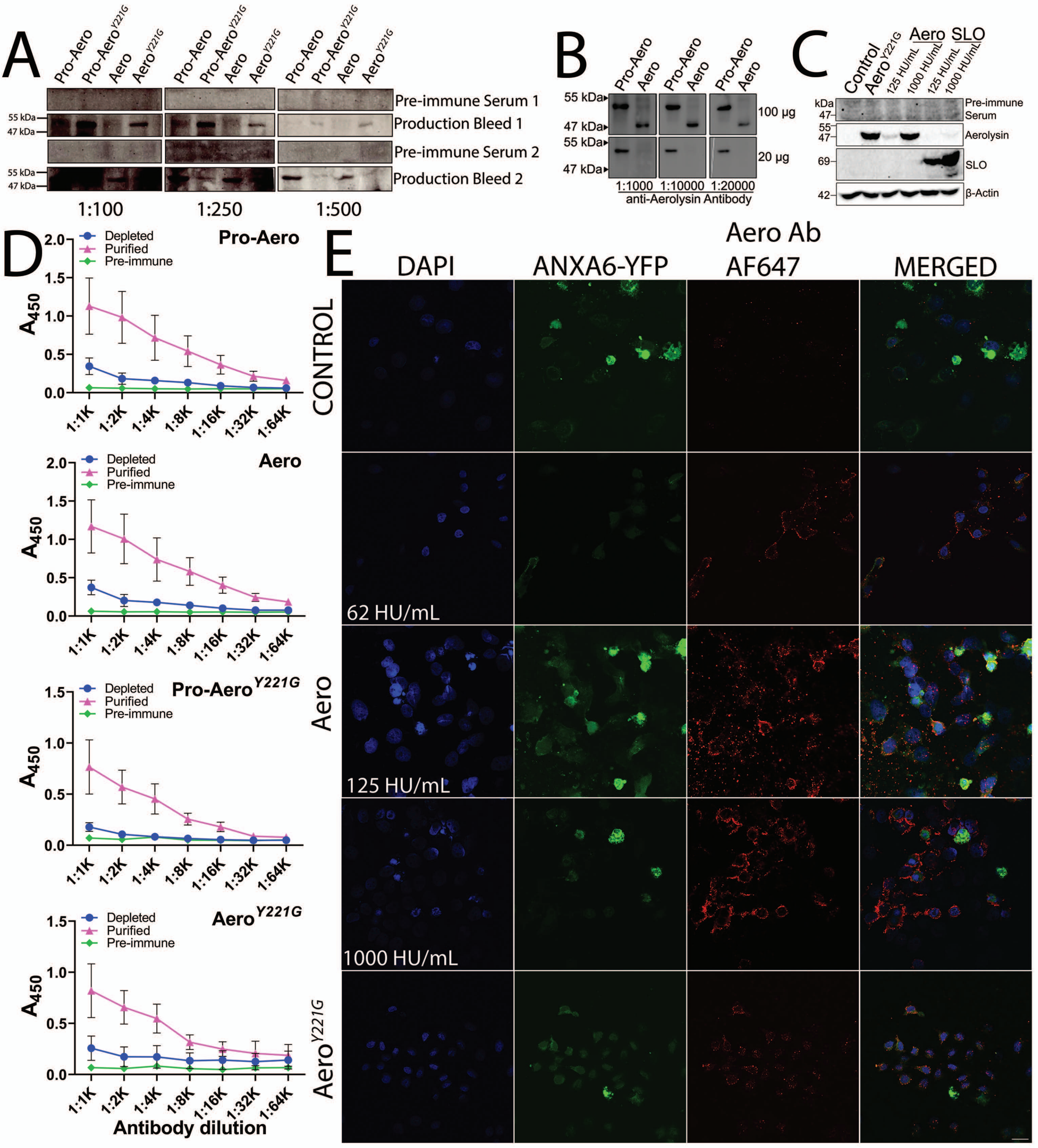

A
Pro-Aero
Pro-Aero Y221G
Aero
Aero Y221G
Pre-immune Serum 1
Production Bleed 1
Pre-immune Serum 2
Production Bleed 2
55 kDa
47 kDa
1:100
1:250
1:500
B
100 µg
20 µg
1:1000
1:10000
1:20000
anti-Aerolysin Antibody
C
Control
125 HU/mL
1000 HU/mL
SLO
kDa
Pre-immune Serum
Aerolysin
β-Actin
69
42
D
Depleted
Purified
Pre-immune
A450
1:1K
1:2K
1:4K
1:8K
1:16K
1:32K
1:64K
Antibody dilution
E
DAPI
ANXA6-YFP
Aero Ab AF647
MERGED
CONTROL
62 HU/mL
1000 HU/mL

### Figure S9

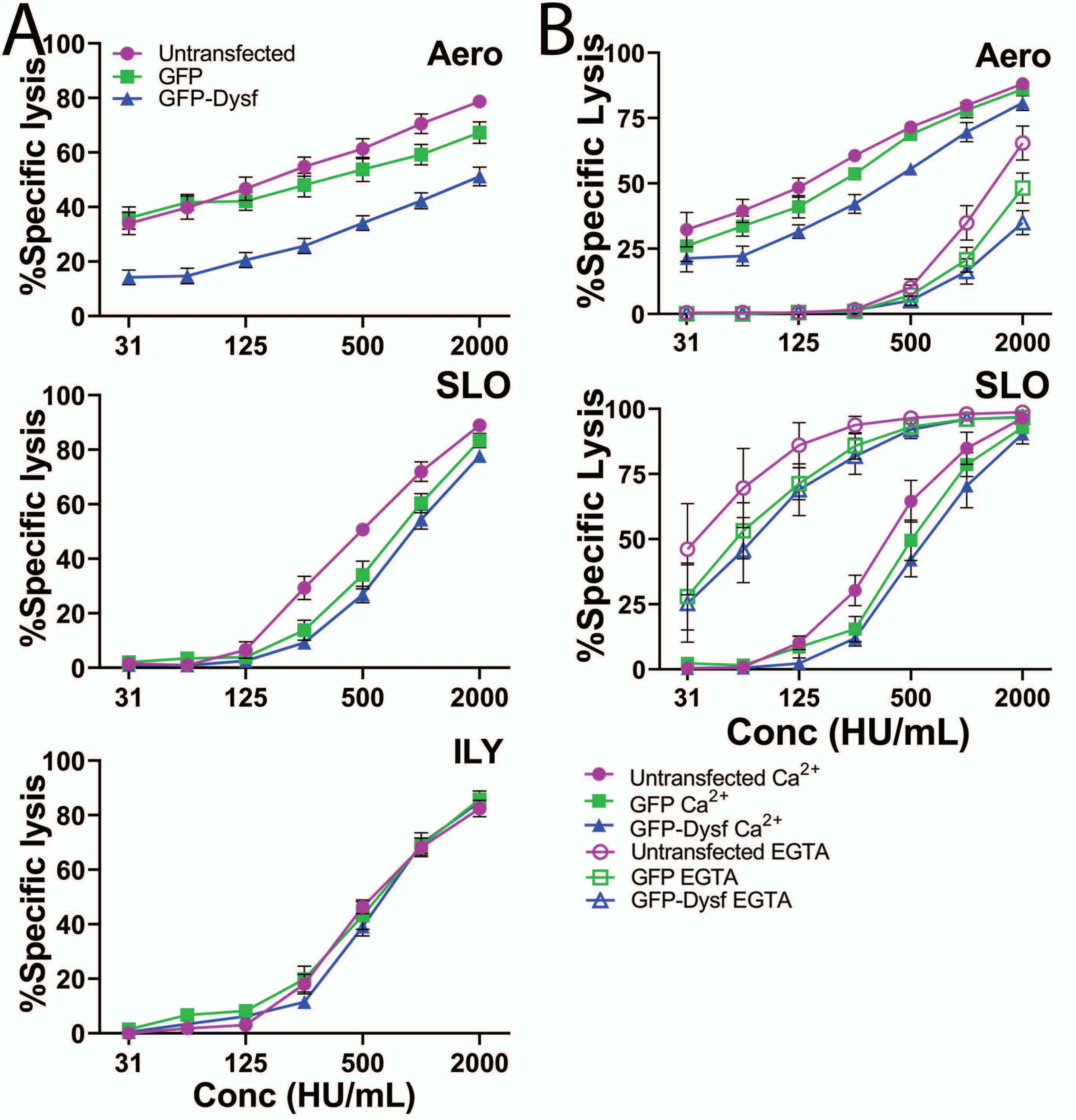
A
B
Aero
SLO
ILY
%Specific lysis
%Specific Lysis
Conc (HU/mL)
31
125
500
2000
Untransfected
GFP
GFP-Dysf
Untransfected Ca2+
GFP Ca2+
GFP-Dysf Ca2+
Untransfected EGTA
GFP EGTA
GFP-Dysf EGTA

### Figure S10

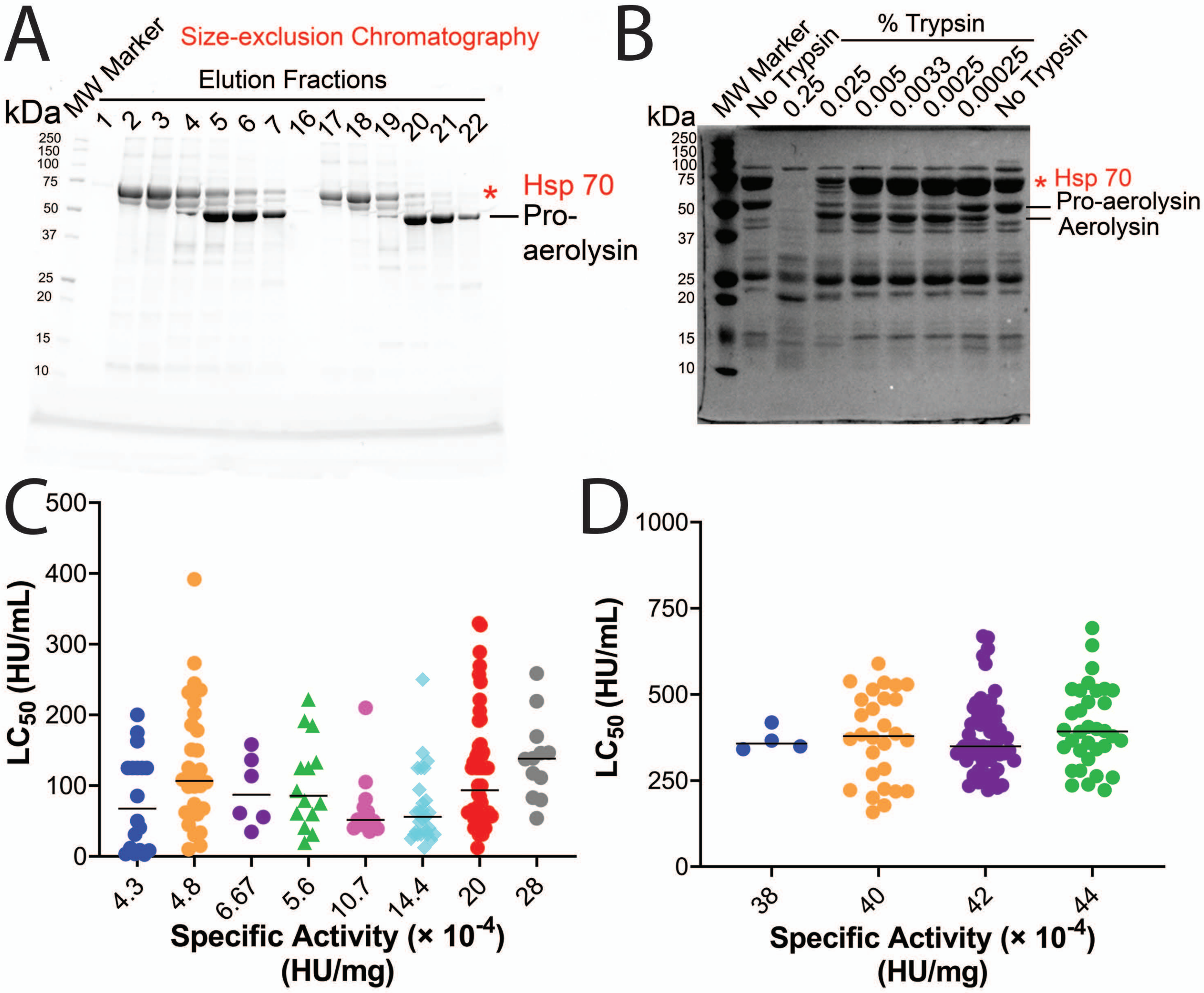
A
MW Marker
Size-exclusion Chromatography
Elution Fractions
kDa
1 2 3 4 5 6 7 16 17 18 19 20 21 22
250
150
100
75
50
37
25
20
15
10
* Hsp 70
Pro-aerolysin
B
MW Marker
No Trypsin
% Trypsin
0.25 0.025 0.005 0.0033 0.0025 0.00025
No Trypsin
kDa
* Hsp 70
Pro-aerolysin
Aerolysin
C
LC50 (HU/mL)
0 100 200 300 400 500
4.3 4.8 6.67 5.6 10.7 14.4 20 28
Specific Activity (× 10-4) (HU/mg)
D
LC50 (HU/mL)
0 250 500 750 1000
38 40 42 44
Specific Activity (× 10-4) (HU/mg)

### Figure S11

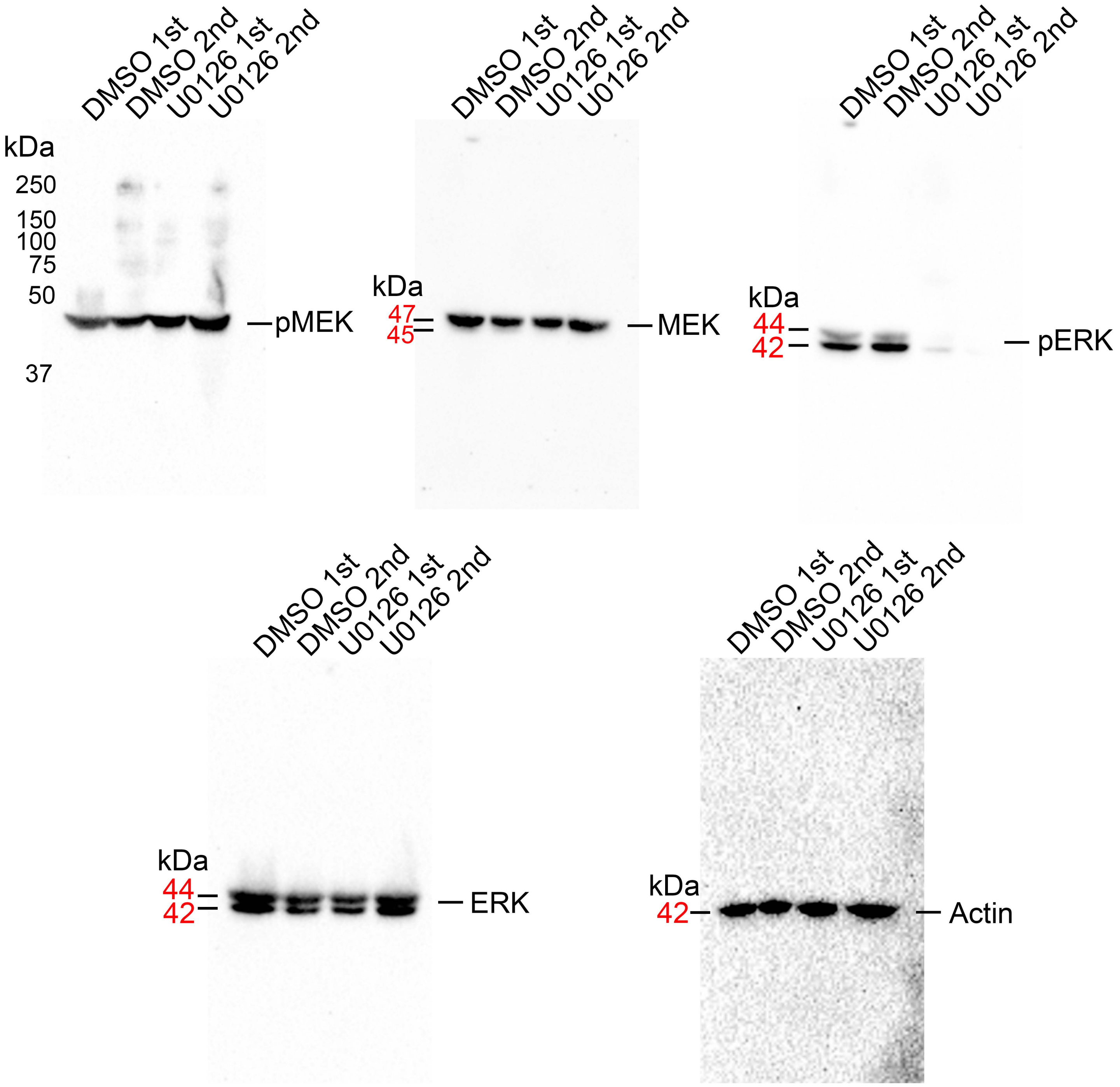

DMSO 1st
DMSO 2nd
U0126 1st
U0126 2nd
kDa
250
150
100
75
50
37
pMEK
47
45
MEK
44
42
pERK
ERK
Actin

### Figure S12

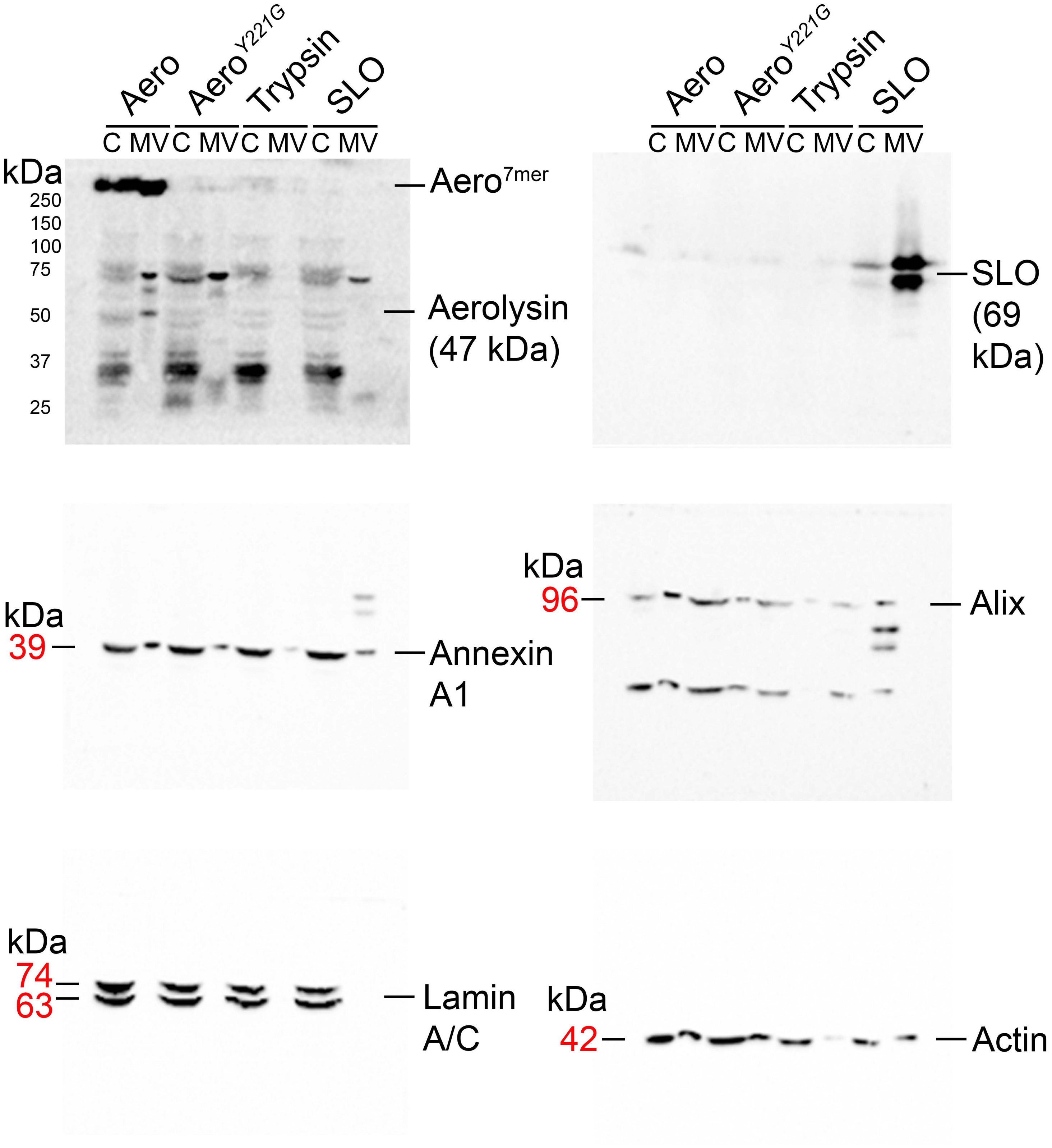

Aero
Aero$^{Y221G}$
Trypsin
SLO
C MV C MV C MV C MV
kDa
250
150
100
75
50
37
25
Aero$^{7mer}$
Aerolysin (47 kDa)
SLO (69 kDa)
kDa
39
Annexin A1
kDa
96
Alix
kDa
74
63
Lamin A/C
kDa
42
Actin
